# Integrated Analysis of Methylome and Transcriptome Following Developmental Atrazine Exposure in Zebrafish Reveals Aberrant Gene-Specific Methylation of Neuroendocrine and Reproductive Pathways

**DOI:** 10.1101/2020.01.28.922179

**Authors:** Chris Bryan, Li Lin, Junkai Xie, Janiel Ahkin Chin Tai, Katharine A. Horzmann, Kyle Wettschurack, Min Zhang, Jennifer Freeman, Chongli Yuan

## Abstract

Atrazine (ATZ) is one of the most commonly used herbicides in the United States. Previous studies have hypothesized the role of ATZ as an endocrine disruptor (EDC), and developmental exposure to ATZ has been shown to lead to behavioral and morphological alterations. Specific epigenetic mechanisms responsible for these alterations, however, are yet to be elucidated. In this study, we exposed zebrafish embryos to 0.3, 3, and 30 ppb (µg/L) of ATZ for 72 hours post fertilization. We performed whole-genome bisulfite sequencing (WGBS) to assess the effects of developmental ATZ exposure on DNA methylation in female fish brains. The number of differentially methylated genes (DMG) increase with increasing dose of treatments. DMGs are enriched in neurological pathways with extensive methylation changes consistently observed in neuroendocrine and reproductive pathways. To assess the effects of DNA methylation on gene expression, we integrated our data with transcriptomic data. Four genes, namely CHD9, FRAS1, PID1, and PCLO, were differentially expressed and methylated in each dose. Overall, this study identifies specific genes and pathways with aberrant methylation and expression following ATZ exposure as targets to elucidate the molecular mechanisms of ATZ toxicity and presents ATZ-induced site-specific DNA methylation as a potential mechanism driving aberrant gene expression.

## INTRODUCTION

Atrazine (6-chloro-N-ehtyl-N-(1-methylethyl)-1,3,5-triazine-2,4-diamine. ATZ) is a widely used agriculture herbicide that controls the growth of broadleaf and weeds. The estimated total annual usage of atrazine is ∼ 76.5 million pounds in the U.S., making it the second most commonly used herbicide in the country(1). ATZ has a relatively high water solubility(2), and thus can be easily traced in surface and ground water by rainfalls and tile drainages(3–5).

ATZ intake from contaminated water supplies has a wide range of adverse health outcomes, including disruption of the hypothalamic-related axis in the brain by inhibiting the luteinizing hormone production and decreasing testosterone production in rats and humans and is thus known to be a potential endocrine disrupter(6–8). Studies have also shown that ATZ exposure can demasculinize and feminize males while disrupting ovarian function in female rats(9). Recent studies also suggest that ATZ disrupts endocrine function by altering the hypothalamic-pituitary-gonadal (HPG) axis(10). For example, a recent work utilizing a female zebrafish animal model has found that ATZ exposure can disrupt the production of luteinizing hormone, follicle stimulating hormone and gonadotropin-releasing hormone, all of which are essential for female reproductive function(11). In addition to its endocrine disruption functions, ATZ exposure has also been associated with several neurological disorders. Studies using a zebrafish animal model have found that ATZ exposure can introduce significant alterations to neurotransmission pathways resulting in significant decreases of serotonin metabolites in female fish(12). Similar observations were made in mouse models, but ATZ exposure was found to primarily impact dopaminergic neurons(13–16). Interestingly, a recent epidemiological study shows that agricultural workers are at high risk of developing depression and anxiety symptoms compared to other occupational groups(17), and the production of serotonin are associated with depression and anxiety in humans(18). Increasing evidence indicates that ATZ-induced neurotoxicity is sex-specific, with females having a higher chance of developing non-motor symptoms(19). Few studies, however, exist using female animal models, and the mechanisms of sex-specific neurotoxicity remain unclear.

The toxic effects of ATZ can persist after removal of exposure source and can be inherited trans-generationally(20), suggesting the existence of underlying molecular “memory” and thus alluding to the involvement of an epigenetic mechanism. Epigenetics, including DNA methylation, histone post-translational modifications, and RNA-assisted regulation(21), refer to reversible and inheritable changes in gene expression without altering the underlying DNA sequence. Among these, the alteration of DNA and histone methylation profiles has been the most well-studied due to its abundance and relative stability compared to other types of epigenetic modifications(22–24). Specifically, a mouse study has found that ATZ exposure can result in deregulation of tissue-specific RNA transcription up to the third generation by inheritable decreases in H3K4me3, a transcription activation marker(25). Similar observations were also made in a rat model(20, 26). Our recent work has demonstrated that ATZ exposure in zebrafish can decrease global DNA methylation levels by inhibiting the activity of maintenance DNMTs and the expression level of DNMT4 and DNMT5, a zebrafish ortholog for DNMT3b(27), but that global methylation in the adult female zebrafish brain was not changed (manuscript in review). There is, however, no genome-wide studies revealing the impact of ATZ exposure on the epigenome in the brain. In this study, we used zebrafish as an animal model and performed whole-genome bisulfite sequencing using brains harvested from 9 month-old fish that were exposed to ATZ only during embryogenesis (1-72 hour post fertilization; hpf) to reveal permanent DNA methylome changes arising from embryonic ATZ exposure. Our results suggest that genes involved in key pathways related to observed behavioral changes resulting from low-dose ATZ, such as neuroendocrine function, are differentially methylated in the brain of zebrafish exposed to ATZ. After integrating the methylation data with transcriptomic data, we suggest the ATZ-induced site-specific DNA methylation as a potential mechanism driving the aberrant expression of neurological genes.

## MATERIAL AND METHODS

### Zebrafish husbandry and treatment

Aliquots of a 10 parts per million (ppm; mg/L) stock solution of technical grade (98.1% purity) ATZ (CAS 1912-24-9; Chem Service, West Chester, PA) were diluted with filtered aquaria water to make exposures of 0.3, 3, and 30 ppb ATZ as previously described, with filtered aquaria water serving as the 0 ppb control(12, 28). Embryos were collected from a breeding colony of AB wild-type zebrafish (Danio rerio) maintained in a Z-Mod System (Aquatic Habitats, Apopka, FL) with a 14:10 light-dark cycle, temperature of 26-28°C, pH 7.0-7.3, and conductivity 470-550 µS. Fish and aquaria were monitored twice daily and fed a mixture of brine shrimp (Artemia franciscana; Artemia International LLC., Fairview, Texas), Zeigler adult zebrafish food (Zeigler Bros Inc., Gardners, PA), and Golden Pearls 500-800 µm (Artemia International LLC., Fairview, Texas). To obtain embryos, adult zebrafish were bred in spawning tanks according to established protocols(29, 30). Embryos were collected at the 4-8 cell stage, randomly sorted into the 0, 0.3, 3, or 30 ppb ATZ exposure groups, and incubated in petri dishes with 20 mL of exposure solution at 28.5°C. At 72 hpf, the larvae were rinsed in filtered aquaria water to end the ATZ exposure and then reared under normal conditions until nine months post fertilization (mpf) (**Fig. S13**) (Supporting Information). At nine mpf, the zebrafish were sexed and three randomly chosen females from each exposure were euthanized via anesthetic overdose with 0.4 mg/ml buffered tricaine-S (ethyl m-amino benzoate methanesulfonate; Western Chemical Inc., Ferndale, WA) and the brains dissected. Female zebrafish were chosen for this study due to the previously reported altered neurotransmission and gene expression changes in female zebrafish brain(12). All protocols were approved by the Purdue University Animal Care and Use Committee and all fish treated humanely with regard to prevention and alleviation of suffering.

### gDNA extraction and bisulfite conversion

gDNA was collected from nine mpf female zebrafish brain following a standard protocol modified for brain samples(27, 31). Zebrafish brains were homogenized in lysis buffer (50 mM Tris, 100 mM EDTA, 100 mM NaCl, 1% SDS, 100 µg/mL Proteinase K) and incubated overnight at 55°C with rocking agitation. The next day, samples were transferred to 1.5 Phase Lock Gel (PLG; QuantaBio, Beverly, MA) tubes, and phenol (phenol-Tris saturated, pH 8; Roche, Indianapolis, IN) and chloroform:isoamyl alcohol (American Bioanalytical, Natick, MA) were added before the samples were centrifuged at room temperature for 5 minutes at 1500 rcf. The upper aqueous phase was transferred to a new PLG tube and the previous steps repeated. The upper aqueous phase was transferred to a 2 ml screw top microcentrifuge tube (Fisher Scientific, Hampton, NH) and 0.1X volume of 3M sodium acetate and 1X volume of isopropanol were added and the tube inverted until DNA aggregation. Samples were incubated at room temperature and then centrifuged at 4°C for 10 minutes at 800 rcf to form a DNA pellet. The pellet was washed in 70% ethanol twice, centrifuging at 4°C for 10 minutes at 800 rcf between washes. After the second wash, the pellet was centrifuged again and dried to remove remaining ethanol. The sample was rehydrated in 1xTLE buffer and incubated overnight at 55°C with rocking agitation. The following morning, the DNA quality and concentration was checked with a NanoDrop ND-1000 Spectrophotometer (Thermo Scientific, Wilmington, DE) and stored at 4°C until further analysis.

For bisulfite conversions, ∼500 ng of gDNA were bisulfite-converted using EZ DNA Methylation-Gold Kit (Zymo Research, CA, USA) following the manufacturer’s protocol. Bisulfite-converted gDNA were eluted to a final concentration of ∼ 100 – 200 ng/μl and stored at −20°C prior to sequencing. The quality of BS-converted samples, such as fragment size, were further examined via a Picochip (Agilent Bioanalyzer, U.S.). The fragment size was found to peak at ∼ 1000 nt as expected.

### Bisulfite Sequencing

Next generation sequencing was performed using NovaSeq (Illumina, US) at Purdue Genomics Core Facility. All sequencing was performed at a sequencing depth of 30x coverage on the 1.5 billion base zebrafish genome. The zebrafish genome (GRCz11) was used as the reference genome.

### Statistical and data analysis

All samples were collected in triplicates. The basic read quality was first verified via summaries produced by the FastQC program (Babraham Institute). Reads were then aligned to the zebrafish reference genome (GRCz11) using Bowtie 2(32), and the read mapping and methylation extraction was performed using Bismark v.0.21.0 (Babraham Institute)(33). A minimum alignment score function (f(x) = 0 + -0.6x, where x is the read length) was used. Reads aligned to identical positions on the genome were removed and bases that have more than 99.9th percentile of coverage in each sample were discarded to account for excessive PCR amplifications. In addition, reads with coverage below 10x were removed to increase the power of statistical tests.

Statistical difference in the global average CpG methylation was determined using a one-way ANOVA and F test followed by Tukey’s HSD post-hoc test (n = 12). A similar test was performed using the average CpG methylation of CpG sites located within 100 bp of the TSS (n = 12).

Differential methylation analysis was performed using the methylKit software package in R(34). First, the genome was tiled with windows of 1000 bp in length and 1000 bp in step-size. Then, a logistic regression model was fitted to predict the log-odds ratio of methylation proportion in each tile using data from all replicates. A sliding linear model (SLIM) method was used to correct P-values to q-values in order to correct for multiple hypothesis testing. Regions with a methylation change greater than 30% and a q value less than 0.01 are considered as differentially methylated regions (DMRs). DMRs were then annotated using the Ensembl GRCz11 zebrafish build via the annotatr software package(35).

Locations of CpG islands, introns, exons, and TSS were determined using the Ensembl GRCz11 build. For all analysis, the promoter region is considered 2000 bps upstream of the coding region due to the lack of annotated promoter sequence in zebrafish genome. Gene ontology and functional enrichment analysis was performed using the PANTHER classification system(36). Zebrafish gene nomenclature was converted to human orthologs using Ensembl TransMap alignments in the UCSC genome browser. Subsequently, Ingenuity Pathway Analysis (IPA) was performed using human orthologs. Only experimentally-validated genes were used for pathway analysis.

Hierarchal clustering and principal component analysis were performed using R version 3.5.1. For hierarchal clustering, the Euclidean distance was used as the distance metric while McQuitty’s method was used as the linkage method.

### Integrated analysis of WGBS with transcriptomic data

The WGBS data was integrated with zebrafish microarray data from our previous study attained from brain tissue of adult female zebrafish aged 6 months with the same embryonic ATZ exposure(12). Overlapping genes were determined using Ingenuity Pathway Analysis (IPA).

## RESULTS

### Initial alignment and quality control

Initial quality control of the sequencing data was performed using FastQC (Babraham Institute), where reads were filtered based on duplicated sequences to ensure that at least 20% of the reads were unique (**Fig. S1**) (Supporting Information). Reads were also filtered by GC content to ensure that the distribution of GC content throughout the genome did not deviate significantly from a theoretical normal distribution (**Fig. S2**) (Supporting Information). Subsequently, reads were aligned to the z11 zebrafish reference genome using Bowtie 2 and Bismark. Alignment rates of reads ranged from 46.4% to 50.6% (**Table S2**) (Supporting Information). **Fig. S3** (Supporting Information) illustrates the frequency of read coverage, which remains consistent between treatments.

### Low dose ATZ exposure results in formation of DMRs at different chromosomal locations

We selected to work with ATZ doses of 0, 0.3, 3 and 30 ppb. The embryonic exposure of these doses is known to result in immediate transcriptomic alterations for genes associated with nervous system development(12). The United States Environmental Protection Agency has implanted a maximum contaminant level (MCL) of 3 ppb (ppb; μg/L) for ATZ levels in drinking water. Brain samples collected from zebrafish 9 mpf with embryonic ATZ exposure were compared here.

We first profiled the type of DNA methylation abundant in zebrafish brain. The results are shown in **Fig. 1**. Throughout all samples, the vast majority of methylated cytosines were in CpG context consistent with other bisulfite sequencing data in zebrafish and mammals(37).

**Figure 1.**
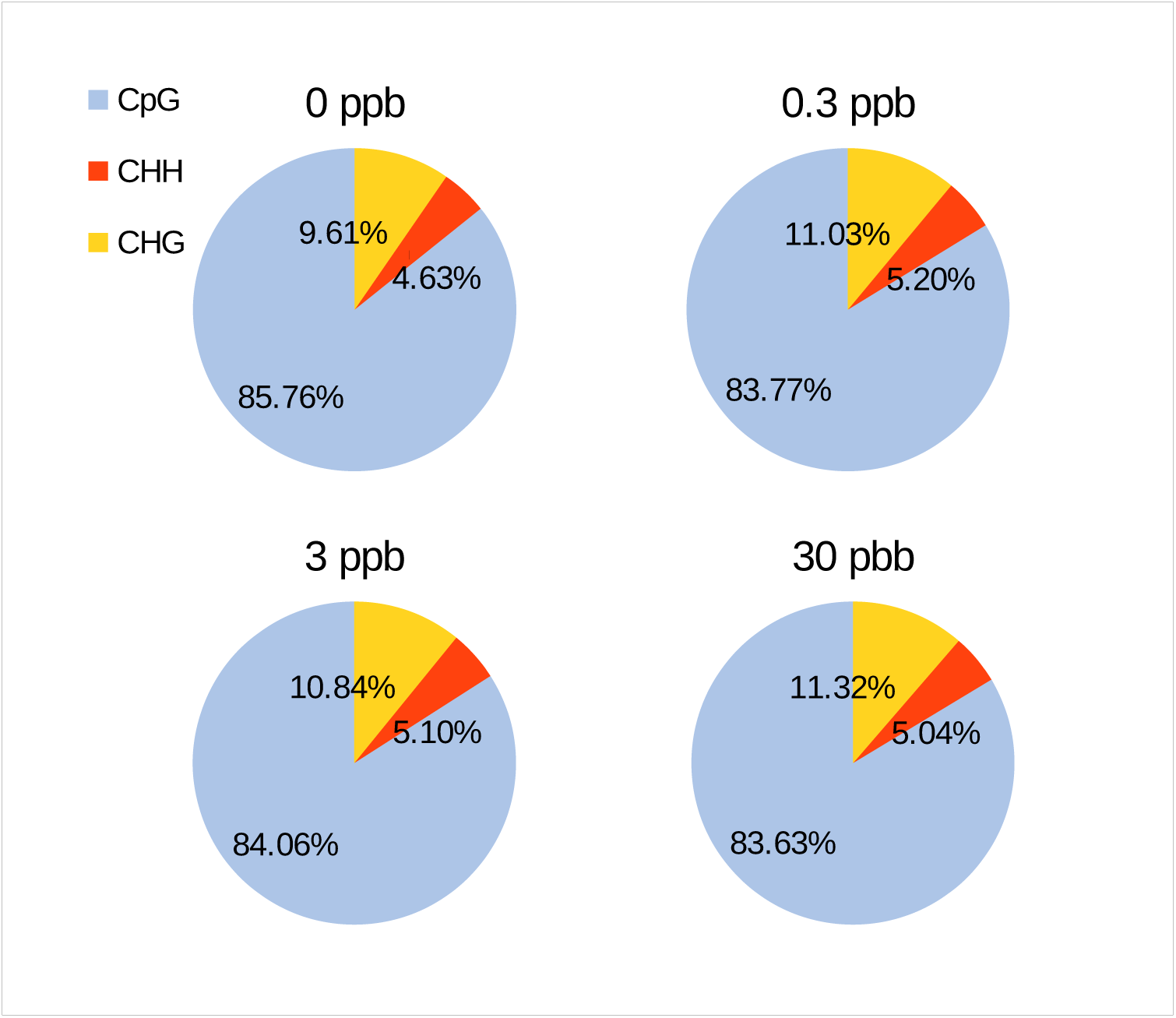
The sequence context of methylated cytosines in the genome at different concentrations of atrazine are compared. Percentages represent the proportion of methylated cytosines found in CpG, CHH, and CHG sequence contexts at 0, 0.3, 3 and 30 ppb ATZ.

CpG methylation was found to be .8074 ± .0043, .8086 ± .0027, .8071 ± .0056 and .8103 ± .0043 for 0, 0.3, 3, and 30 ppb treated zebrafish brains respectively, as shown in **Table S3** (Supporting Information). There is, however, no statistically significant difference in CpG methylation percentage among treatments (*p* = 0.371). **Fig. 2A** depicts the distribution of CpG methylation throughout the genome. The proportion of CpG methylation was mainly concentrated in three distinct clusters: CpG sites with all cytosines methylated, CpG sites with ∼ 75 - 95% of cytosines methylated, and CpG sites with no cytosines methylated. Different treatments exhibited similar distributions of CpG methylation with the aforementioned pattern. **Fig. 2B** and **C** illustrate the average CpG methylation per gene within exons and promoter regions respectively. In line with **Fig. 2A**, the genome-wide distribution of methylation was very similar among treatments both for the exon and promoter regions. Most promoter regions exhibit average methylation between 75% and 90%, and the frequency of CpG methylation at proportions below 75% is similar. In exons, an even larger proportion of genes contain methylation between 75% and 90%, and the proportion of genes with 50% methylation or less is lower than that of the promoter regions.

**Figure 2.**
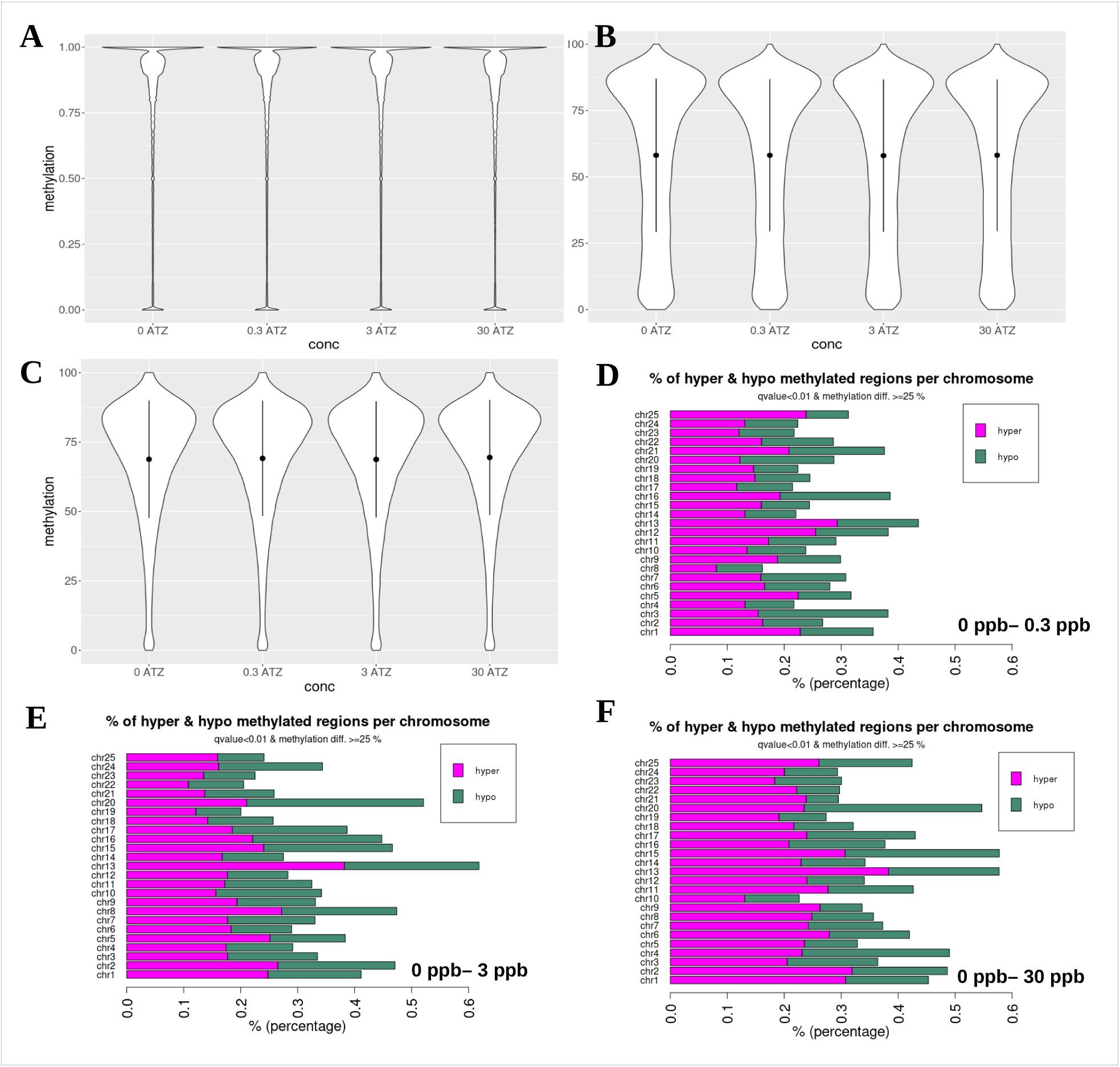
Genome-wide Profile of CpG Methylation. A. Violin plot of the proportion of methylated cytosines at each CpG site in the genome. B-C. Violin plot of the proportion of methylated cytosines at promoter regions (1000 bp upstream of TSS) and exon regions respectively. D-F. Percentage of hyper- and hypo-methylated regions per chromosome in samples treated with 0.3 ppb (D), 3 ppb (E), and 30 ppb (F) of ATZ.

We then identified specific regions in the genome that exhibited significantly different levels of methylation between treatments, also known as differentially methylated regions (DMR), and grouped them based on their chromosomal locations. We categorized DMRs into hyper- and hypo-methylated regions, where hypermethylated DMRs exhibit increased methylation level after exposure to ATZ while hypomethylated DMRs exhibit decreased level of methylation. Among comparisons to untreated zebrafish samples, we identified 1587, 1826, and 2455 genes containing hypermethylated DMRs and 1186, 1442, and 1226 genes containing hypomethylated DMRs in 0.3 ppb, 3 ppb, and 30 ppb, respectively. **Fig. 2D-F** demonstrate the percentage of hypermethylated and hypomethylated regions per chromosome in 0.3 (**D**), 3 (**E**), and 30 (**F**) ppb ATZ-treated samples as compared to untreated zebrafish brains. **Fig. S4** (Supporting Information) further illustrates changes induced by ATZ treatment at different chromosomal locations using a heatmap. **Fig. S5** (Supporting Information) further defines DMRs by comparing samples treated with varying doses of ATZ.

### Functional Features of Identified DMRs

To better assess the sequence context and genetic location of DMRs, we annotated genes within DMRs using the GRCz11 reference genome. Genes containing at least one DMR in either the promoter or gene body are considered as a differentially methylated gene (DMG). **Fig. 3A** illustrates the locations of DMR between gDNA treated with 0.3 and 0 ppb of ATZ. The largest fraction of DMRs (∼ 38%) were found in introns, seconded by intergenic regions (∼ 37%). ∼ 14 and 11% of DMR was found to be within exon and promoter regions, respectively. These functional regions contain different CpG contexts. Specifically, intron, intergenic, exon and promoter regions each consist of ∼ 45.37, 40.60, 6.94 and 7.04% of CpG sites. We can thus correct the abundance of DMRs by accounting for their CpG context and found that the proportion of DMR within intron, intergenic, exon and promoter is 0.83, 0.93, 1.97, and 1.28 times as of the proportion of all CpG sites, respectively. The fraction of DMR within these four selected categories were found to be in a close accordance with each other independent of ATZ doses as shown in **Fig. S6** (Supporting Information).

**Figure 3.**
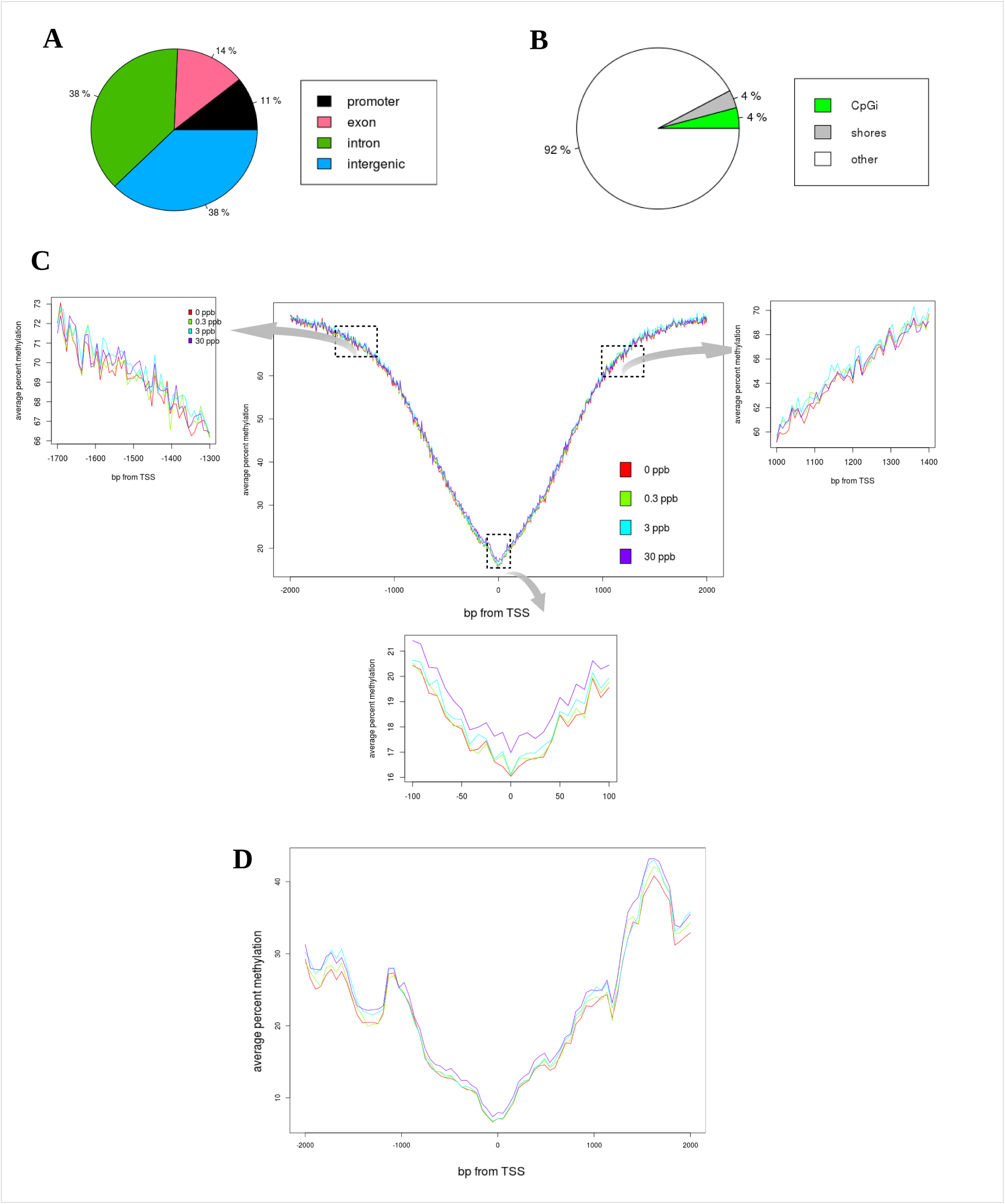
Genomic Location of Differentially Methylated Regions. A. Proportion of DMR located in exons, introns, promoters (1000 kb upstream), and intergenic regions for 0 - 0.3 ATZ comparison. B. Proportion of DMR located in CpGi, CpG shores and other CpG-containing sequences for 0 - 0.3 ATZ comparison. C. Average methylation percentage 2000 bp up- and downstream of the transcription start site (TSS). Zoom-in views of selected regions (highlighted in dotted squares) are also provided. D. Average methylation percentage of genes containing CpGi (-2000 to 2000 bp of TSS).

We then compared the CpG context of DMR as shown in **Fig. 3B**. 4% of DMRs (between 0 and 0.3 ppb) are located in CpG island (CpGi) and another 4% of DMRs are located in CpG flanking regions, also known as CpG shores. The rest, and majority, of DMRs are located within other sequence contexts. CpGi and shores were identified in our study via the CpG island track from the UCSC Genome Browser. The fraction of DMRs located within different CpG contexts is found to be similar when comparing samples treated with different ATZ concentrations as shown in **Fig. S7** (Supporting Information). Within the zebrafish genome, 1.96% of CpG sites are located in CpGi while 3.34% are located in CpG shores. Thus, the proportion of DMR located in CpGi and CpG shores is over-represented compared to the rest of CpG sites.

DNA methylation in promoter regions are critically important for regulating gene activity. We thus plotted average methylation percentage in genetic regions immediately adjacent to the transcriptional starting site (TSS) with 8 base-pair resolution as shown in **Figs. 3C**. Within approximately ∼ 100 bp up- and down-stream to TSS, samples exposed to 30 ppb ATZ exhibit the highest level of methylation, with the difference most distinctive at the TSS itself (*p* = .0000536) (see **Fig. 3C** bottom). In this same range, samples exposed to 0.3 and 3 ppb ATZ exhibit similar levels of methylation, but both have higher methylation levels than untreated samples at most locations. The difference is only significant, however, between the TSS methylation levels of 30 ppb and other concentrations. Furthermore, we clustered promoter regions based on their respective CpG contexts and illustrated the methylation distribution near TSS of genes containing promoters with CpGi or CpG shore as shown in **Fig. 3D** **and Fig. S8** (Supporting Information), respectively. Samples exposed to 0.3 and 3 ppb ATZ exhibit nearly identical methylation levels to untreated samples near the TSS but deviate significantly in distinct spikes upstream of the TSS. Interestingly, we noticed a significant ^me^CpG drop in all treatment and control groups near both ∼1.3 and ∼1.7 kb upstream of the TSS, and the difference among treatments is most distinctive within this region. Genes with promoters containing CpG shores (**Fig. S8** (Supporting Information)) exhibit similar methylation changes as other genetic regions as shown in **Fig. 3C**.

Subsequently, we performed principal component and hierarchal clustering analysis in order to elucidate heterogeneity in DNA methylation patterns among treatment groups and between replicates of the same treatment. **Fig. 4A** illustrates the separation of all measurements mapped to a PC space. The PC space was determined via scaling of all CpG sites throughout the genome. Untreated samples have very similar methylation patterns and are in a well-defined cluster within the PC space. Replicates exposed to 0.3 and 3 ppb ATZ have more diverse DNA methylation features, with some replicates closely resembling the methylation features of untreated samples and others developing distinctive methylation features. The methylation features are most distinct within replicates exposed to 30 ppb ATZ. Similar analysis was performed using methylation of promoter and exon sequences as shown in **Figs. S9A and C** (Supporting Information). To further validate this observation/hypothesis, hierarchal clustering (HC) of samples based on methylation patterns was performed. Compared to PCA, HC provides a more quantitative metric of similarity between replicates of differing concentrations. **Fig. 4B** shows the clustering results of all replicates. There are 3 main clusters found in the dendrogram: one consisting of 3ATZa and c, 0.3ATZa and b, 30ATZb, and 0ATZa-c, another consisting of 3ATZb and 0.3ATZc, and a third consisting of 30ATZa and c. The trend observed in HC is consistent with our findings from PCA using genome-wide CpG methylation features, and supports the hypothesis that response to ATZ between replicates is dose-dependent. A similar analysis was performed using the methylation pattern of promoter and exon regions as shown in **Fig. S9** (Supporting Information).

**Figure 4.**
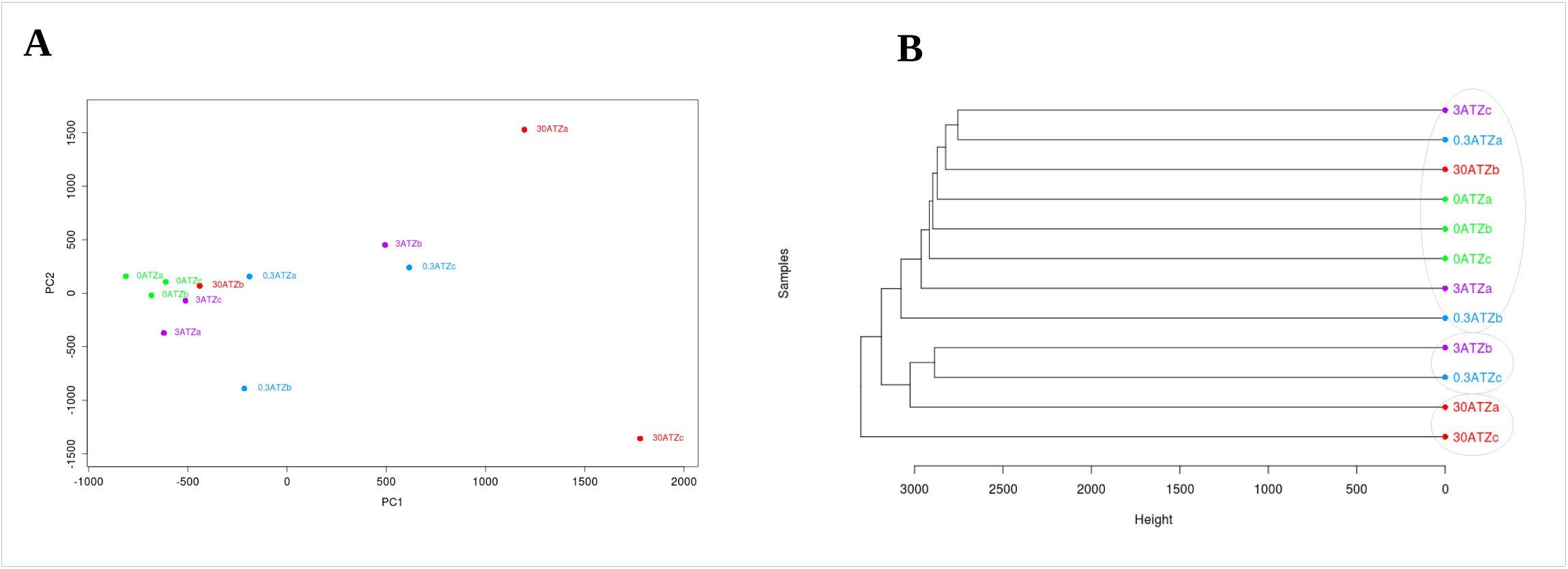
Cluster Analysis based on CpG Methylation Profiles A. Principal Component Analysis (PCA) of samples constructed with all CpG sites. B. Hierarchal clustering of samples based on centroid distance of methylation percentage at all CpG sites.

### Low-dose atrazine exposure results in aberrant methylation in key neurogenic pathways

Gene Ontology (GO) analysis was performed to identify the function and genetic pathways of regions that were differentially methylated. **Figs. 5A** and **5B** illustrate the number of hypo- and hyper-DMG, respectively, between treatment and control group. **Figs. S10A** and **S10B** (Supporting Information) summarizes the number of DMGs in promoter and gene body for each treatment group compared to the control. The number of DMGs increased with increasing ATZ concentration in almost all cases. **Figs. 5C** and **5D** show the representative % of hypo- and hyper- methylated DMGs clustered by protein functions when comparing treated groups to the control. PANTHER protein classification database was used for gene classification. Among them, transcription factor, enzyme modulator, hydrolase and nucleic acid binding protein were consistently found most affected by ATZ treatments in both hyper and hypo-methylated cases.

**Figure 5.**
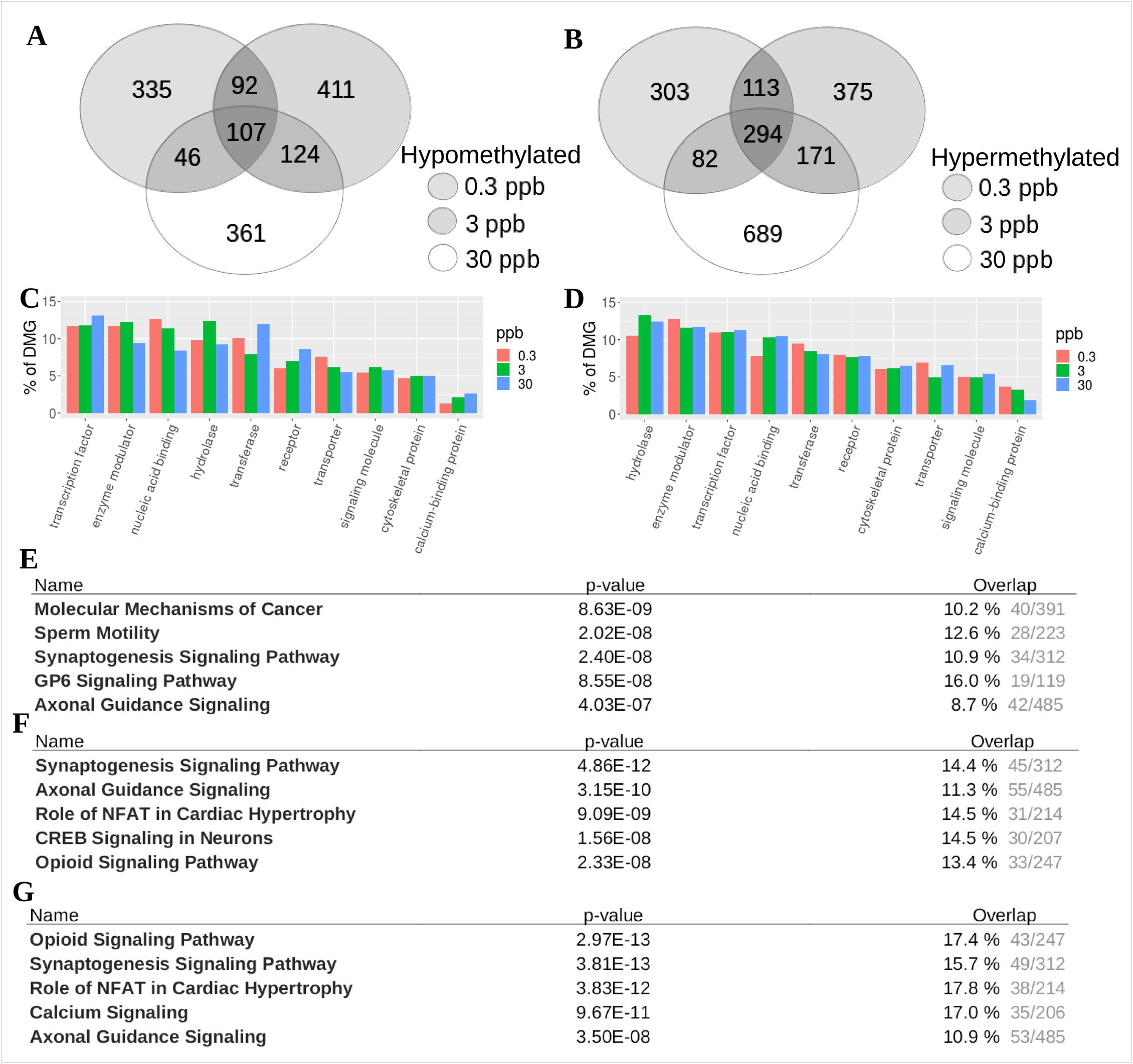
Gene Ontology (GO) Analysis of Differentially Methylated Genes. A-B. Number of (A) hypomethylated and (B) hypermethylated DMG in each comparison group. C-D. Protein function of (C) hypo- and (D) hyper-methylated DMG comparing to un-treated control. E-G. Highest represented IPA pathways of DMG in (E) 0 – 0.3 ppb, (F) 0 - 3 ppb, and (G) 0 – 30 ppb ATZ comparison. Overlap was reported as the proption of affected genes within the pathway.

**Figs. 5E-G** illustrates the top pathways from IPA using DMRs identified by comparing samples exposed to control. Among top 5 pathways sorted by p-value, the neuronal pathways of Synaptogenesis Signaling and Axonal Guidance are present in each comparison group. Top diseases and upstream regulators affected are listed in **Tables S4** and **S5** (Supporting Information), respectively. Cancer, organismal injury and abnormalities, endocrine system disorders and gastrointestinal diseases are identified in all exposed groups comparing to the untreated control. Interestingly, cancer-related regulators, i.e., TP53 and PHIP, are primarily identified in low ATZ doses, while levodopa, the precursor of dopamine, is identified as an upstream regulator in 3 and 30 ppb ATZ treated samples.

We selectively focused on 3 neuronal pathways here, namely GABA receptor signaling, long-term synaptic depression and gonadotropin-releasing hormone signaling. All three pathways were significantly enriched in all concentrations of ATZ. **Figs. 6A-C** demonstrate DMG within the GABA Receptor, Long-term Synaptic Depression, and Gonadotropin-Releasing Hormone Signaling pathways, respectively for brain samples exposed to 0.3 and 30 ppb of ATZ. Equivalent results for samples exposed to 3 ppb of ATZ can be found in **Fig. S11** (Supporting Information). The number of DMGs within these selected pathways increase as ATZ concentration increases. The additional DMGs found in 30 ppb are highlighted in dotted circles in **Fig. 6**.

**Figure 6.**
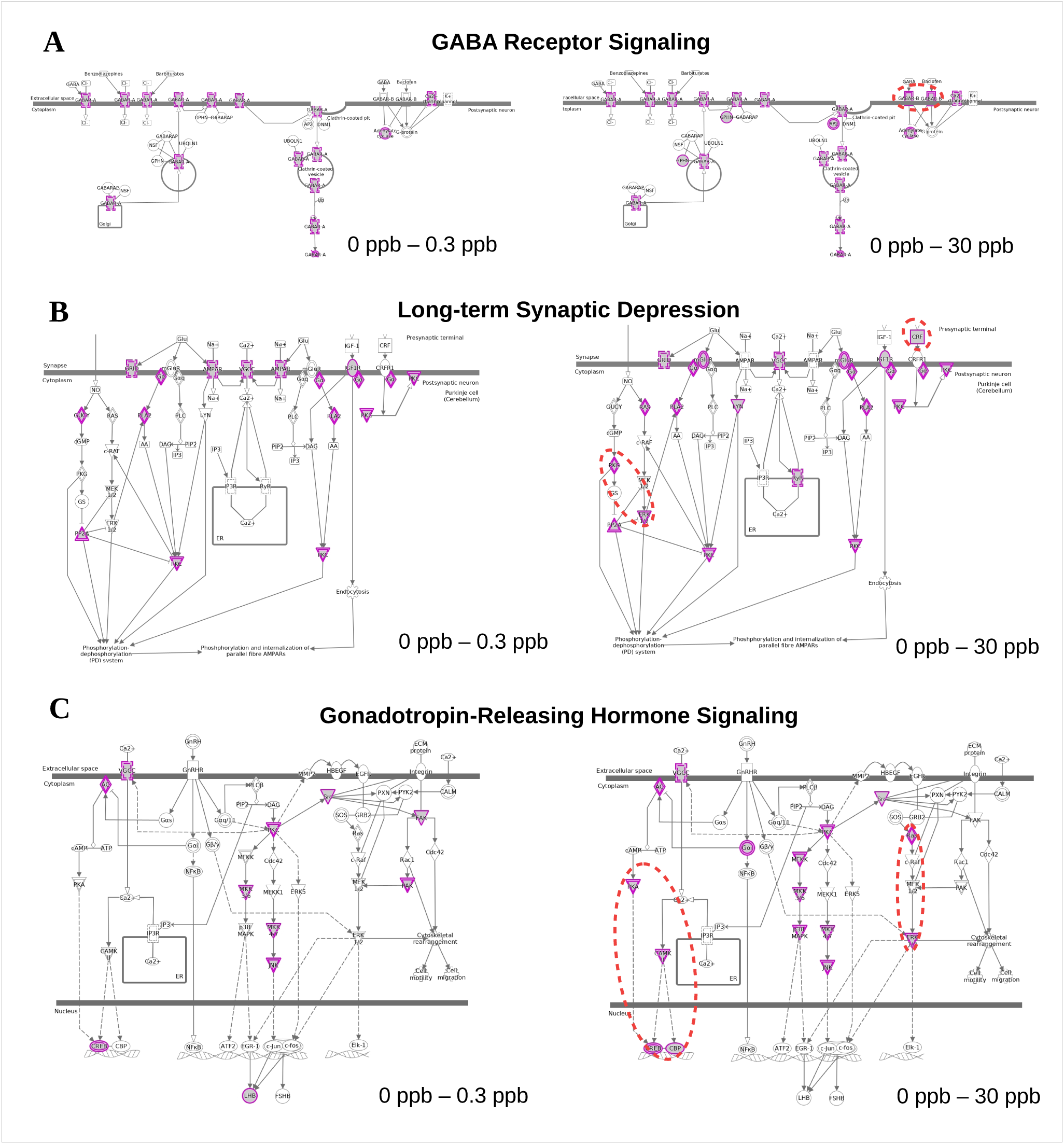
IPA Analysis of Differentially Methylated Genes in Neurogenic Pathways. DMG (marked in purple) within (A) the Synaptogenesis Signaling pathway, (B) Long-Term Synaptic Depression pathway, and (C) Gonadotropin-Releasing Hormone Signaling pathway found in 0- 0.3 ppb (left) and 0 – 30 ppb (right) comparison. DMGs that uniquely emerge at high dose of ATZ (30 ppb) were highlighted by red dotted circles.

### Integrated analysis of bisulfite-seq with transcriptomic data

In order to elucidate the potential correlation of methylation patterns with gene expression resulting from ATZ exposure, we integrated data from bisulfite sequencing (this study) with gene expression data from microarray collected using zebrafish (6 mpf) with embryonic ATZ exposure(12). We noticed the difference in our selected time-points, but the microarray data in (12) was collected from zebrafish with identical exposure to ATZ as we described in this work and thus provide a reasonable comparison point. **Fig. 7A** shows the number of differentially expressed genes adapted from literature(12) with the largest number of differentially expressed genes observed in 0.3 ppb. **Fig. 7B** integrates both datasets and illustrates genes that were both differentially expressed and methylated. The largest number of common genes was observed in 0.3 ppb treated groups. Four genes were found both differentially expressed and methylated common in each treatment group.

**Figure 7.**
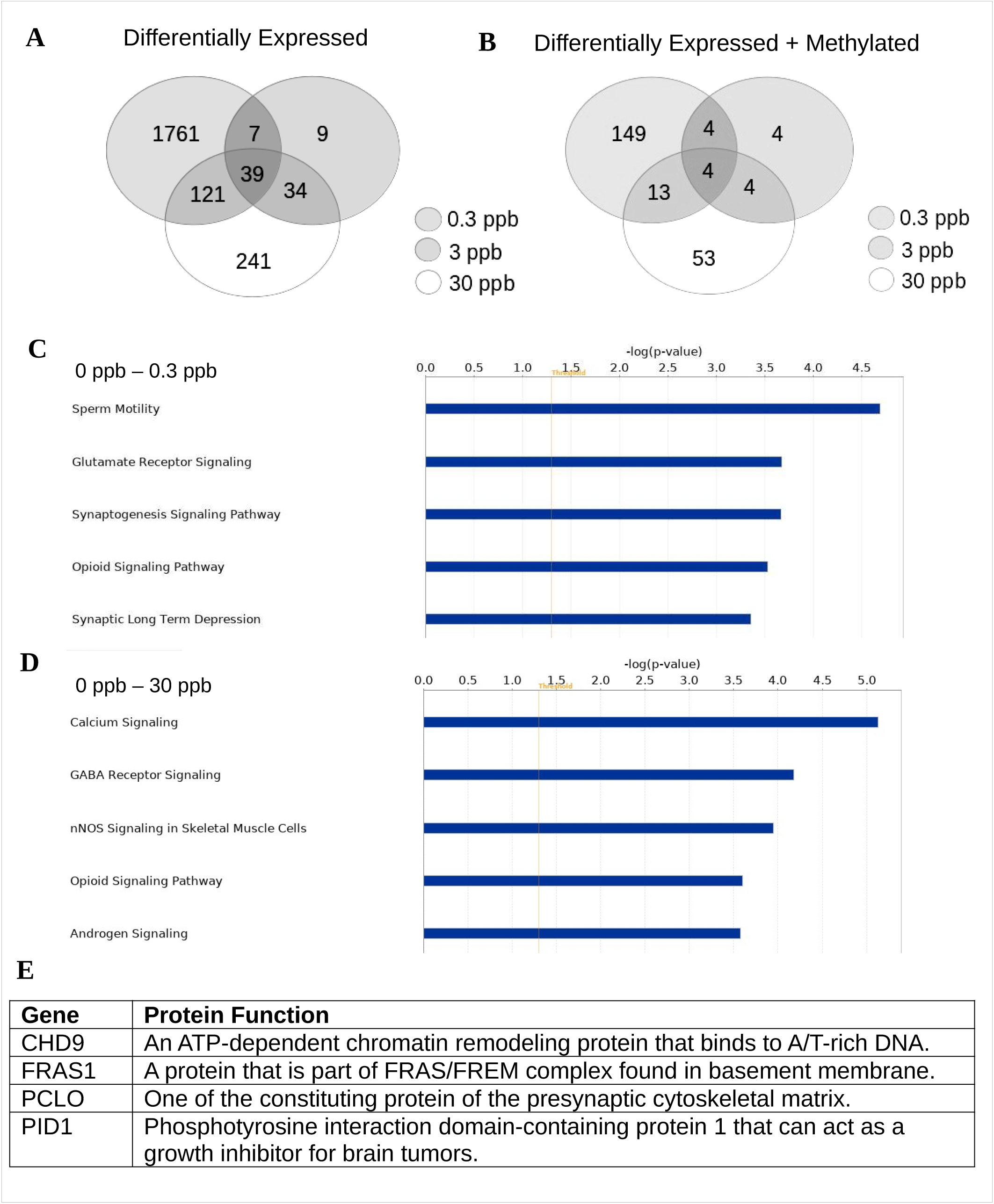
Integrated Analysis of Differentially Methylated and Expressed Genes. A. Venn diagram representing overlapping differentially expressed genes between different concentrations of ATZ adapted from (12). B. Venn diagram representing overlapping genes between differentially expressed genes and differentially methylated genes, C-D. Top five IPA pathways of overlapping genes (both differentially expressed and methylated) in (C) 0 – 0.3 ppb and (D) 0– 30 ppb comparison. E. Description of biological functions of human orthologs to 4 common genes that are differentially expressed and methylated in each comparison.

Pathways of commons genes for 0.3 and 30 ppb of ATZ treatment compared to untreated control can be found in **Figs. 7C** and **D** (Supporting Information). Results form 3 ppb are not shown here due to the small number of genes that are both differentially expressed and methylated. The protein functions of 4 common genes identified in all treatment doses are summarized in **Fig. 7E**.

Furthermore, we plotted the fold change in gene expression level (Y-axis) versus the percentage of methylation change (X-axis) of all genes with DMRs located at either promoter, exon or intron regions as shown in **Fig. 8**. **Figs. 8A-C** illustrates common genes with both methylation and expression level changes after treating with 0.3, 3 and 30 ppb of ATZ. Each gene was also color-coded based on its expression level. **Figs. 8D-F** illustrate genes with DMR located in promoter regions. Fold change in expression level and methylation was correlated using a linear fitting as shown in **Figs. 8D-F**. A negative correlation was found at all treatment doses. A similar analysis was carried using genes with DMRs located in exons or introns as shown in **Fig. S12** (Supporting Information).

**Figure 8.**
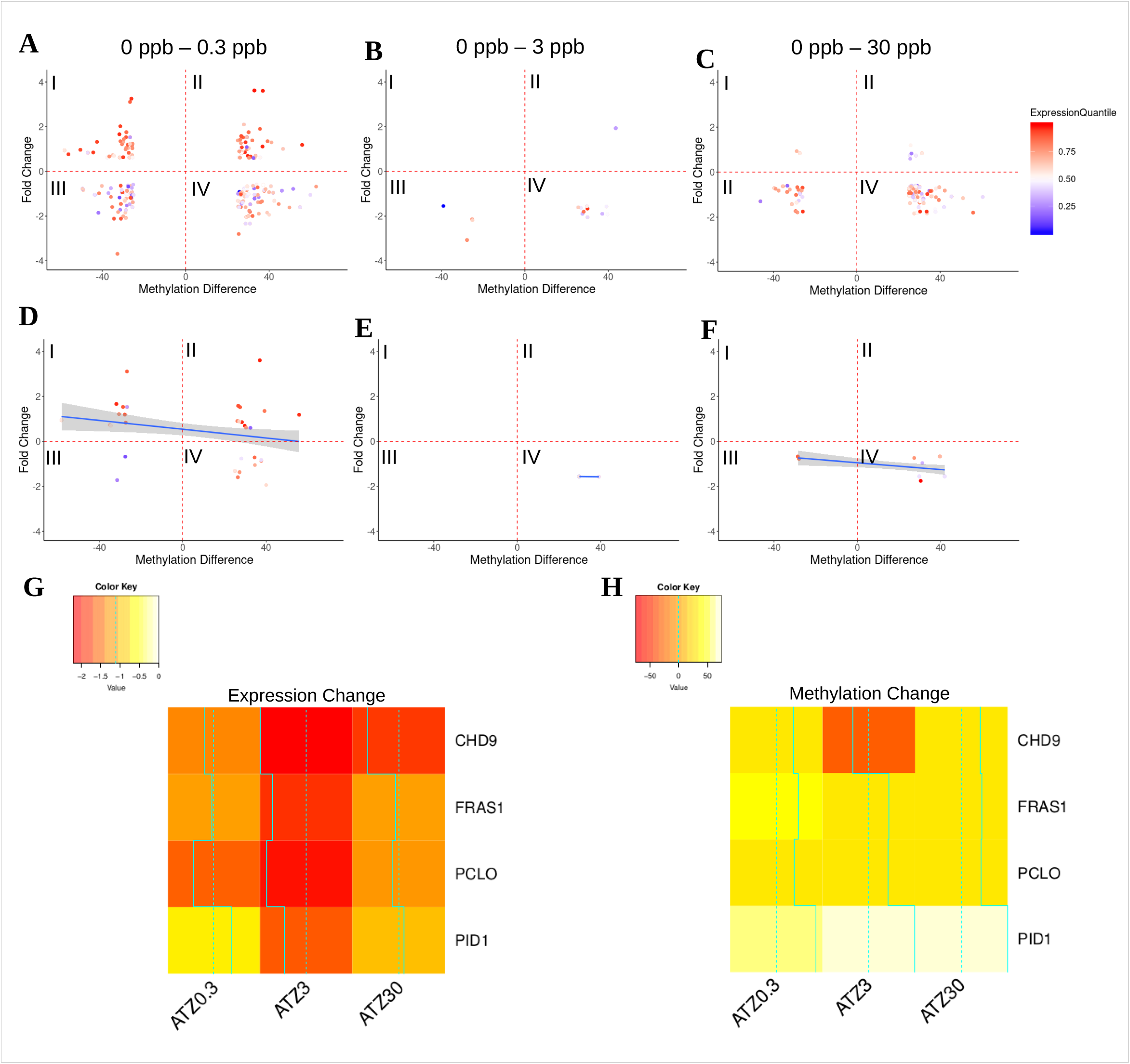
Correlation Between DNA Methylation and Gene Expression, Grouped by Expression Level. Scatterplots of differentially methylated and expressed genes with methylation difference plotted (x-axis) against fold change in protein expression level (y-axis) for DMGs identified in (A) 0 - 0.3 ppb, (B) 0 - 3 ppb and (C) 0 ppb - 30 ppb comparisons. The expression level of DMGs were color coded. A similar set of scatterplots was prepared for DMGs containing differentially methylated promoter regions identified in (D) 0 – 0.3 ppb, (E) 0 – 3 ppb and (F) and 0 - 30 ppb comparison. Heatmap comparing (G) gene expression and (H) methylation levels of four genes both differentially methylated and expressed across all ATZ concentrations.

We then focus on the four genes that were both differentially expressed and methylated in each treatment group. **Fig. 8G** illustrates the fold change in expression level of these genes while **Fig. 8H** illustrates their change in DNA methylation after exposing to ATZ of varying doses.

## DISCUSSION

### Embryonic ATZ exposure results in persistent changes in DNA methylation pattern

The United States Environmental Protection Agency has set a maximum action level of 3 ppb for atrazine in drinking water. Drinking water with atrazine concentrations exceeding 3 ppb, however, are often reported in various regions, particularly Midwestern region of the U.S. (38). We thus selected to work with ATZ concentrations of 0.3, 3 and 30 ppb in this work. Brain samples were collected from zebrafish 9 mpf with embryonic exposure to ATZ for 72 hrs. Our WGBS results suggest that all samples have ∼ 81 % cytosine methylation with majority of methylation (∼ 84%) existing in a CpG context. Methylation in CHG and CHH sites are also observed, but with significantly lower abundance. Compared to CpG methylation, the biological role of CHG and CHH methylation are not as well established; we thus focus on CpG methylation on our follow-up analysis.

The global distribution of methylated CpG sites within different functional regions and CpG contexts does not seem to be significantly altered after exposure to ATZ as shown in **Figs. 2A-C** and **3A-B**. A significant fraction of genes was found to contain hypo- or hypermethylated genes (DMGs) after embryonic ATZ exposure across all chromosomes compared to the untreated control as shown in **Figs. 2D-F** and **S4-5** (Supporting Information). The amount of DMGs increase with increasing ATZ concentrations in most chromosomal locations. Specifically, the amount of hypermethylated regions increased from 0.3 to 3 ppb ATZ in 19/25 chromosomes (72%) and from 3 to 30 ppb ATZ in 21/25 chromosomes (84%). In each concentration of ATZ, the largest proportion of DMR was consistently found in chromosome 13, while chromosome 8 and 2 exhibited the largest fold change between low and high concentrations of ATZ (1.92 and 2.82 fold respectively). The amount of hypomethylated regions increased from 0.3 to 3 ppb ATZ in 11/25 chromosomes (44%) and from 3 to 30 ppb ATZ in 7/25 chromosomes (28%). In total, there were more hypermethylated DMR than hypomethylated DMR in each concentration, and hypermethylated DMR also exhibited the largest dose-dependent effect, suggesting that ATZ may play a more significant role in the addition of methyl groups.

The increase in both hypo- and hyper-methylated regions are likely to cancel out each others’ effect on changes in CpG methylation on a genome-wide scale, and subsequently result in no alterations of global CpG methylation. The number of DMRs increase with increasing ATZ doses suggesting a potential close to linear dose-dependence. The largest proportion of DMRs (hyper + hypo) were identified in chromosome 13 and chromosome 20. Collectively, our results suggest that ATZ exposure selectively alter DNA methylation based on the chromosomal locations, while global modification level and distribution remains minimally perturbed. Similar observations were made in zebrafish embryos using other EDCs including BPA and dioxin, both of which were found to elicit no significant changes in global CpG methylation levels, but on specific gene sets (37, 39).

Functionally, CpG methylation are selectively enriched in exon and promoter regions consistent with their expected roles in modulating gene expression(40). Accounting for CpG contents, only a small fraction of DMRs was found in CpGi. This observation can be potentially explained by the small number of CpGi in the zebrafish genome. The amount of CpGi in zebrafish was found to be ∼ 1/10 of what has been reported in other fish types, i.e., stickleback (Gasterosteus aculeatus) and tetraodon (Tetraodon nigroviridis) (41). Compared to other known EDCs, i.e., BPA, the percentage of differentially methylated CpG islands was significantly higher in zebrafish exposed to ATZ. For example, the number of genes containing methylated CpG islands in zebrafish exposed to 0.3 ppb ATZ [111] was almost twice as high as that found for zebrafish exposed to 10 μM of BPA [64] (37).

A clustering analysis was carried out accounting for CpG methylation features of all genes (**Fig. 4**) and in selected functional regions, namely promoter and exon regions (**Fig. S9** (Supporting Information)). Interestingly, our results suggest that although the detailed methylation pattern vary among replicates, samples treated with no or low-dose (0.3 and 3ppb) of ATZ are self-similar among the treatment groups. High dose of ATZ (30 ppb) treatments result in more heterogeneous features in DNA methylation among the treated group. Similar conclusions can be made using CpG methylation features of exons but not promoter regions. Collectively, our results suggest that ATZ exposure can alter methylation patterns, but the changes may be heterogeneous with increased heterogeneity associated with higher ATZ concentrations. This conclusion, however, can be further strengthened from having a larger sample size.

Our results thus unequivocally suggest that embryonic ATZ exposure can result in DNA methylome change in a dose-dependent manner. The changes are more enriched in certain chromosomal locations including chromosome 13 and 20. Increased methylome heterogeneity is also observed as a result of ATZ exposure.

### Low-dose atrazine exposure results in aberrant methylation in key neurogenic pathways

GO analysis was used to identify the functional role of DMGs and their affiliated pathways. When comparing treated to untreated control group, enzyme modulator, hydrolase, nucleic acid binding, and transcription factor proteins are found to consist the largest fraction of DMGs (**Fig. 5C-D**). Interestingly, the number of nucleic acid binding protein increases with higher doses of ATZ in hypermethylated genes, while decreasing with higher dose in hypomethylated genes.

Among pathways identified in IPA, synaptogenesis and axonal guidance signaling pathways are the top identified pathways among all treatment groups. Opioid signaling pathway is identified in samples treated with 3 and 30 ppb of ATZ. Synaptogenesis signaling pathway plays a critical role in the formation of synapses. Synaptogenesis primarily occurs during developmental stages and is almost completed silenced during adulthood via DNA methylation(42). Dysregulation of this pathway is affiliated with the pathogenesis of autism(42). Axonal guidance signaling pathways guide axons to their targets to form synaptic connections vital for nerve growth in brains and other sensory tissues, e.g., retina of eyes(43). Abnormal DNA methylation of genes in this pathway as been associated with non-syndromic high myopia(44). Opioid signaling pathway mainly consists of opioid receptors, i.e. G protein-coupled receptor (GPCR), and is involved in multiple physiological processes including pain signaling, reproduction and immunological response(45–47). Hypermethylation of opioid receptors has been associated with psychosis and AD (48, 49). These top affiliated pathways thus provide strong evidence that ATZ exposure can result in significant CpG pattern changes in critical neurogenic pathways which provide the molecular foundation for observed neuro-behavior changes that has been reported in ATZ exposed animal models(50).

The wide range of biological functions and the vast number of constituent genes in the aforementioned neuronal pathways made it difficult to identify and interpret dose-dependent changes in DNA methylation of neuroendocrine and neurotransmitter signaling pathways. Therefore, we focused on three selected pathways, namely GABA receptor signaling, long-term synaptic depression, and gonadotropin-releasing hormone signaling pathways. These pathways were selected because they were significantly enriched in all comparison groups. They involve axonal and synaptic genes and thus can represent top pathways identified in our IPA analysis (**Figs. 5E-G**). Among them, 4, 6 and 6 genes were found to consistently exhibit different CpG methylation levels in GABA receptor signaling, long-term synaptic depression and gonadotropin-releasing hormone signaling pathway, respectively, after treating with ATZ. The number of genes affected in each pathway also increase with increasing dose of ATZ.

GABA is the most abundant inhibitory neurotransmitter found in mammalian central nervous systems. GABA receptors are transmitter-gated ion channels. The amount of GABA receptor on surface can determine the strength of inhibitory synapses. Changes in GABA receptor expression is known to be regulated by DNA methylation (51) and affiliated with various psychiatric diseases, including schizophrenia-like symptoms and post-traumatic stress disorder(52, 53). Exposure to chlorotriazine herbicide has been previously reported to cause changes in GABA_A_ activity(54). Interestingly, at low dose of ATZ (0.3 ppb), only GABA_A_ receptors and GABA-gated Cl-channels are differentially methylated. At high dose of ATZ (30 pbb), GABA_B_, a G protein coupled receptor activated by baclofen, also becomes differentially methylated suggesting more extensive impacts on GABA receptor signaling. GABA has also been show to interact with the hypothalamus–pituitary–gonadal (HPG) axis, as it regulates GnRH and LH release(55).

Long-term synaptic depression accounts for an activity-dependent reduction in synaptic transmission efficiency of neurons and has been associated with motor learning(56). In all doses of ATZ, differential methylation within synaptic receptor proteins of this pathway were observed, including IGFR-1, GRID, and AMPAR. In 30 ppb ATZ, methylation of metabotropic glutamate receptor (mGluR) and corticotropin-releasing factor (CRF) was observed. DNA methylation and other epigenetic mechanism has been implicated in modulating synaptic plasticity(57). Abnormal methylation of identified genes have been broadly affiliated with an array of neurodegenerative and psychiatric diseases. For example, abnormal methylation in group III metabotropic glutamate receptor pathway genes have been associated with Alzheimer’s Disease, Parkinson’s Disease, stress and anxiety. Studies have shown that activation of mGluR7 is associated with increased levels of stress hormone corticosterone and corticotrophin which are known to trigger stress- and anxiety-like symptoms(58), in line with observed social behavior of zebrafish following ATZ exposure(59). Changes in locomotor activity has been previously reported in both male and female rats exposed to ATZ and attributed to disruptions in dopaminergic systems(14, 60). Disrupted long-term synaptic depression can thus be a plausible underlying mechanism accounting for changes in locomotion of animals exposed to ATZ.

Gonadotropin-releasing hormone (GnRH) signaling pathway is a central regulator for the reproductive system and thus aligns with the suspected role of ATZ as an EDC. Alteration in this pathway can result in behavioral and phenotypic changes, such as female-specific alteration of GABA levels (see previous paragraph) and endocrinal disruption of the hypothalamus–pituitary–gonadal (HPG) axis, both of which has been previously identified in animals with atrazine exposure(6, 12). Exposure to 0.3 ppb ATZ was observed to increase methylation levels of central protein kinases within the GnRH pathway as well as gonadotropic luteinizing hormone (LH) and CREB. LH promotes ovulation and develops the corpus luteum in females, and low-dose ATZ exposure in rats has been shown to disrupt LH pre-ovulatory surge(61). At higher dose of ATZ (i.e., 30 ppb), in addition to the central kinase genes of the pathway, the cAMP/PKA pathway and their downstream targets CREB and CBP are also differentially methylated. CREB and ERK1/2 (also differentially methylated) are critical regulators of progesterone production, and ATZ exposure has been shown in rats to induce enhanced cAMP/PKA contributing to premature luteinization(62).

Cancer was identified as the top affected disease in all doses by IPA, consistent with previous studies indicating the association between embryonic ATZ exposure and altered expression of genes and protein abundance in cancer-related pathways(28, 50). TP53, a well-studied tumor suppressor gene implicated in the onset of cancer(63), was the top predicted upstream regulator in 0.3 ppb and 3 ppb treated samples. Taken together, these results support the hypothesis that ATZ has carcinogenic effects, and present aberrant DNA methylation pattern as a plausible instigator of carcinogenesis.

### Integrated methylome and gene expression analysis reveals potential methylation driven transcriptome changes after exposure to ATZ

Common genes are identified via integrated methylation and gene expression analysis as shown in **Fig. 7**. Since only a few genes were found to be differentially expressed in samples treated with 3 ppb of ATZ, we focused on 0.3 and 30 ppb here. Among genes both differentially expressed and methylated, the top enriched pathways consist of a large number of neurogenic pathways such as synaptogensis signaling, opioid signaling, and calcium signaling similar as we observed in our differential methylation analysis (**Figs. 5E-F**). Furthermore, GnRH signaling, and Long-term Synaptic Depression are enriched in common genes within both 0.3 ppb and 30 ppb, and GABA receptor signaling is enriched in common genes within 30 ppb. Taken as a whole, these results suggest DNA methylation as a driver contributing to transcriptomic aberration within neuroendocrine pathways. Dose-dependent mechanisms of methylation and gene expression within the GABA receptor warrants further study due to its role as a regulator of endocrine hormone signaling.

To further reveal the potential correlation between methylation level and gene expression, we constructed methylation-expression change plots as shown in **Fig. 8** accounting for the expression level of genes in untreated samples. Among all treatment doses, highly expressed genes (+ 50%) seem to be more affected by ATZ (70.4, 68.7 and 69.7% for exposure to 0.3, 3 and 30 ppb of ATZ). When treated with 0.3 ppb of ATZ, genes are almost evenly distributed among all four quadrants. Genes with DMRs in promoter regions are populated in I and IV quadrants, suggesting a negative correlation between ^me^CpG levels with gene expression, consistent with their known biological role. As ATZ concentration increases, more genes are found to locate at III and IV quadrant, suggesting decreased expression level after exposure to ATZ. Specifically, at 30 ppb, most genes affected are enriched in III and IV quadrant with decreased expression levels, among which genes with DMR in promoter regions predominantly exist in IV quadrant. Collectively, ATZ exposure at increased concentrations seems to primarily contribute to down-regulating gene expression. The regulation can be partially attributed to increased methylation level in promoter regions based on our correlation analysis.

A closer look of methylation features near TSS further supports our hypothesis, as DNA methylation increases in promoter regions with increasing ATZ concentrations (**Fig. 3C** and **D**) independent of CpG contexts. Interestingly, we also noticed a significant ^me^CpG increase in all treatment and control groups near both ∼1.3 and ∼1.7 kb upstream of the TSS in genes containing CpGi, and the difference among treatments is most distinctive within this region. This region is typically populated with super-enhancers or distal promoter elements(64). Increase in ^me^CpG within these functional elements may result in decreased expression levels of genes not immediately following the sequence and can thus potentially account for other down-regulated genes found after exposure to ATZ.

A total of four common genes were identified that exhibit consistent methylation and expression level changes across all selected ATZ concentrations. All 4 genes were down-regulated and exhibited a significant increase in ^me^CpG, with the exception of CHD9 methylation in samples exposed to 0.3 ppb ATZ. Taken together with results from the previous paragraph, this observation suggest that the observed CpG methylation changes may be driving the expression changes in these genes.

We identified human orthologs of these four genes with their primarily functions listed in **Fig.7E**. Specifically, CHD9 gene encodes for chromodomain helicase DNA binding protein 9 belonging to a CHD family of proteins governing the access of cellular machineries to DNA. CHD9, also known as chromatin-related mesenchymal modulator, specifically promotes the expression of osteocalcin to promote bone development(65, 66). Altered expression of CHD9 has been observed in gastric and colorectal cancers(67). PCLO is involved in synaptic transmission and plays a significant role in monoaminergic neurotransmission in brain. Mis-regulation of PCLO has been associated with major depressive disorders(68). Interestingly, depression-like neurobehavioral alterations have also been reported in ATZ-exposed zebrafish (69). Fraser extracellular matrix complex subunit 1 (FRAS1), regulates epidermal-basement membrane adhesion and organogenesis during development(70). Mutations and abnormal expression of FRAS1 can result in Fraser Syndrome 1, and differential methylation on FRAS1 promotor region has been observed in tamoxifen resistant and fulvestrant resistant breast cancers(71). Phosphotyrosine interaction domain containing 1 (PID1) is a retention adapter protein that regulates activity of the endocytic receptor LDL receptor-related protein 1 (LRP1). Low expression level of PID1 is observed in glioblastomas and associated with poor prognosis (72).

### Conclusions

Taken together, our results suggest that embryonic ATZ exposure can result in persistent methylation changes at specific chromosomal locations while the global methylation level and distribution remain unaffected. The number of differentially methylated regions/genes increase with higher concentrations of ATZ. ATZ treatment primarily affects methylation features of neurogenic genes associated with key pathways, such as GABA receptor signaling, long-term synaptic depression and gonadotropin-releasing hormone signaling pathways consistent with previous behavior studies using zebrafish. Integrated DNA methylation and gene expression analysis reveals that ATZ exposure primarily down-regulate gene expression driven by increased DNA methylation level suggesting a potential underlying mechanism of physiological alterations following developmental ATZ exposure. In particular, the genes CHD9, FRAS1, PID1, and PLCO are promising targets for further study in their role in the developmental toxicology of ATZ due to their consistent epigenetic silencing. Furthermore, the magnitude of ^me^CpG alteration is dependent on dose. Although small doses of ATZ were sufficient to elicit significant alterations in ^me^CpG of neurologic pathways, exposure to higher doses of ATZ substantially increases the number of altered genes within the same pathways, suggesting more profound effects at a higher dose.

## AVAILABILITY

Raw and processed sequencing data are available through GEO Accession number GSE144154.

