## Supporting Information for "Integrated Analysis of Methylome and Transcriptome Following Developmental Atrazine Exposure in Zebrafish Reveals Aberrant Gene-Specific Methylation of Neuroendocrine and Reproductive Pathways"

### Table of Contents


| Supporting Tables | pg. |
|---|---|
| **Table S1.** Summary statistics of sample gDNA, describing the number of reads, total nucleotides, and percentage of sequences lost as a result of next-generation sequencing for each replicate. | S-4 |
| **Table S2.** Mapping rate of samples to zebrafish z11 reference genome using Bowtie 2 in the Bismark alignment suite. Efficiency is defined as the proportion of reads successfully mapped to a region in the reference genome. | S-5 |
| **Table S3.** Total number of cytosines located in CpG context in each sample and the percentage of methylated cytosines in CpG context. | S-6 |
| **Table S4.** Top diseases identified via human orthologs of DMG found in (**A**) 0.3 ppb, (**B**) 3 ppb and (**C**) 30 ppb treated sampled identified via IPA. P-value ranges was defined using Fisher's exact test. | S-7 |
| **Table S5**. Top predicted upstream regulators of human orthologs of DMG found in (**A**) 0.3 ppb, (**B**) 3 ppb, and (**C**) 30 ppb treated samples identified via IPA. | S-8 |
| **Supporting Figures**<br><br>**Figure S1.** Percentage of duplicated sequences of sample gDNA for each replicate performed using FastQC. | S-9 |
| **Figure S2.** GC content of sample gDNA compared with theoretical normal distribution for each replicate performed using FastQC. | S-10 |
| **Figure S3.** Histograms of sample coverage demonstrating the frequency of length of reads within each dose of ATZ in $\log_{10}$ of read coverage per base. | S-11 |
| **Figure S4.** Heatmap comparing the average methylation percentage of all genes. Color scale represents the average proportion of methylated cytosines at CpG throughout the gene, with each bar representing a unique gene. From innermost to outermost ring: 0 ppb ATZ, 0.3 ppb ATZ, 3 ppb ATZ, 30 ppb ATZ. | S-12 |
| **Figure S5. A.** DMR per chromosome in analysis comparing 0.3 ppb ATZ and 3 ppb ATZ, **B.** DMR per chromosome in analysis comparing 3 ppb ATZ and 30 ppb ATZ, **C.** DMR per chromosome in analysis comparing 0.3 ppb ATZ and 30 ppb ATZ. | S-13 |
| **Figure S6.** Proportion of DMR located in exons, introns, promoters (1000 kb upstream), and intergenic regions for 0 ATZ- 3 ppb ATZ comparison, 0 ATZ- 30 ppb ATZ comparison, 0.3 ATZ- 3 ppb ATZ comparison, 0.3 ATZ- 30 ppb ATZ comparison, and 3 ATZ- 30 ppb ATZ comparison. | S-14 |
| **Figure S7.** Proportion of DMR located in CpGi and CpG shores for 0 ATZ- 3 ppb ATZ comparison, 0 ATZ- 30 ppb ATZ comparison, 0.3 ATZ- 3 ppb ATZ comparison, 0.3 ATZ- 30 ppb ATZ comparison, and 3 ATZ- 30 ppb ATZ comparison. | S-15-16 |
| **Figure S8.** Average methylation percentage in DMGs containing CpG shores (-2000 to +2000 bp of TSS) | S-17 |
| **Figure S9. A.** PCA of replicates using CpG methylation of promoter, **B.** Hierarchal clustering of replicates using CpG methylation of promoter, **C.** PCA of replicates using CpG methylation of exon, **D.** Hierarchal clustering of replicates using CpG methylation of exon. | S-18-19 |

| | |
|---|---|
| **Figure S10. A.** Number of genes with differentially methylated promoter in each comparison group. **B.** Number of genes with differentially methylated gene body in each comparison group. | S-20 |
| **Figure S11. A – C.** DMG from 0 ppb – 3 ppb comparison (marked in purple) within the GABA Receptor signaling pathway (**A**), Long-Term Synaptic Depression pathway (**B**), and Gonadotropin-Releasing Hormone Signaling pathway (**C**). | S-21-22 |
| **Figure S12. A – C**. Scatterplots of differentially methylated and expressed genes with methylation difference on x-axis and expression level fold change on y-axis for genes containing differentially methylated exons identified in (**A**) 0 - 0.3 ppb, (**B**) 0 - 3 ppb and (**C**) and 0 - 30 ppb comparisons. Similar plots were made for common genes with differentially methylated intron sequences as shown for genes identified in (**D**) 0 - 0.3 ppb, (**E**), 0 - 3 ppb and (**F**) and 0 - 30 ppb comparisons. | S-23 |
| **Figure S13.** Illustration of procedure for husbandry and treatment of zebrafish embryos. | S-24 |

**Table S1.** Summary statistics of sample gDNA, describing the number of reads, total nucleotides, and percentage of sequences lost as a result of next-generation sequencing for each replicate.

| Replicate | Dose (ppb ATZ) | Total reads | Bases | %seqs lost | %bases lost |
|---|---|---|---|---|---|
| 1 | 0 | 486,481,238 | 66,181,743,403 | 1 | 11 |
| 2 | 0 | 493,320,648 | 67,567,933,989 | 1 | 11 |
| 3 | 0 | 412,952,992 | 54,733,922,905 | 1 | 13 |
| 1 | 0.3 | 623,065,336 | 86,912,938,597 | 1 | 8 |
| 2 | 0.3 | 388,565,462 | 52,591,825,699 | 1 | 11 |
| 3 | 0.3 | 515,814,254 | 70,514,571,875 | 1 | 10 |
| 1 | 3 | 491,618,516 | 69,101,089,988 | 1 | 7 |
| 2 | 3 | 422,840,874 | 58,904,625,902 | 1 | 8 |
| 3 | 3 | 520,167,440 | 71,133,833,726 | 1 | 10 |
| 1 | 30 | 539,576,838 | 75,661,361,730 | 1 | 8 |
| 2 | 30 | 553,374,426 | 75,331,877,280 | 1 | 11 |
| 3 | 30 | 396,846,486 | 52,603,117,497 | 1 | 13 |

**Table S2.** Mapping rate of samples to zebrafish z11 reference genome using Bowtie 2 in the Bismark alignment suite. Efficiency is defined as the proportion of reads successfully mapped to a region in the reference genome.

| **Concentration ATZ, ppb** | **Mapping Efficiency** |
|---|---|
| 0 | 46.4% |
| 0.3 | 50.6% |
| 3 | 49.8% |
| 30 | 47.8% |

**Table S3.** Total number of cytosines located in CpG context in each sample and the percentage of methylated cytosines in CpG context.

| Replicate | Dose | Methylated CpG | Total CpG | % of methylated CpG |
|---|---|---|---|---|
| 1 | 0 | 397025847 | 97462457 | 80.29 |
| 2 | 0 | 397162010 | 94390337 | 80.79 |
| 3 | 0 | 387694452 | 90027825 | 81.15 |
| 1 | 0.3 | 630221786 | 149529328 | 80.82 |
| 2 | 0.3 | 380012474 | 91356877 | 80.61 |
| 3 | 0.3 | 500157577 | 116198380 | 81.14 |
| 1 | 3 | 552046244 | 132861252 | 80.60 |
| 2 | 3 | 443290261 | 101840804 | 81.31 |
| 3 | 3 | 486408691 | 120015579 | 80.21 |
| 1 | 30 | 550928972 | 130619740 | 80.83 |
| 2 | 30 | 504172621 | 120392247 | 80.72 |
| 3 | 30 | 416670553 | 94370607 | 81.53 |

**Table S4.** Top diseases identified via human orthologs of DMG found in (**A**) 0.3 ppb, (**B**) 3 ppb and (**C**) 30 ppb treated sampled identified via IPA. P-value ranges was defined using Fisher's exact test

**A.** 0 ppb – 0.3 ppb

| Name | p-value range | # Molecules |
|---|---|---|
| Cancer | 2.20E-04 - 8.60E-52 | 818 |
| Organismal Injury and Abnormalities | 2.20E-04 - 8.60E-52 | 822 |
| Endocrine System Disorders | 6.92E-05 - 5.80E-51 | 729 |
| Gastrointestinal Disease | 1.76E-04 - 4.18E-41 | 734 |
| Reproductive System Disease | 3.23E-05 - 1.67E-31 | 571 |

**B.** 0 ppb – 3 ppb

| Name | p-value range | # Molecules |
|---|---|---|
| Cancer | 6.71E-05 - 3.54E-59 | 977 |
| Endocrine System Disorders | 4.16E-05 - 3.54E-59 | 873 |
| Organismal Injury and Abnormalities | 6.79E-05 - 3.54E-59 | 987 |
| Gastrointestinal Disease | 4.16E-05 - 5.26E-53 | 884 |
| Dermatological Diseases and Conditions | 1.37E-05 - 1.69E-39 | 647 |

**C.** 0 ppb – 30 ppb

| Name | p-value range | # Molecules |
|---|---|---|
| Cancer | 1.29E-05 - 2.91E-68 | 1062 |
| Organismal Injury and Abnormalities | 1.32E-05 - 2.91E-68 | 1070 |
| Gastrointestinal Disease | 2.69E-06 - 2.74E-67 | 981 |
| Endocrine System Disorders | 2.91E-09 - 3.81E-60 | 944 |
| Reproductive System Disease | 1.15E-05 - 1.28E-39 | 741 |

**Table S5.** Top predicted upstream regulators of human orthologs of DMG found in (**A**) 0.3 ppb, (**B**) 3 ppb, and (**C**) 30 ppb treated samples identified via IPA.

**A**. 0 ppb – 0.3 ppb

| Name | p-value |
|---|---|
| SOX11 | 9.48E-07 |
| TP53 | 3.14E-06 |
| PHIP | 6.00E-05 |
| RPA1 | 8.61E-05 |
| SYVN1 | 8.70E-05 |

**B.** 0 ppb – 3 ppb

| Name | p-value |
|---|---|
| TP53 | 2.20E-06 |
| levodopa | 2.78E-05 |
| topotecan | 5.10E-05 |
| APC | 5.67E-05 |
| ingenol mebutate | 6.72E-05 |

**C.** 0 ppb – 30 ppb

| Name | p-value |
|---|---|
| HTT | 2.39E-08 |
| levodopa | 2.92E-08 |
| HDAC4 | 3.26E-07 |
| Sos | 4.53E-07 |
| ERG | 7.60E-07 |

**Figure S1.** Percentage of duplicated sequences of sample gDNA for each replicate performed using FastQC.

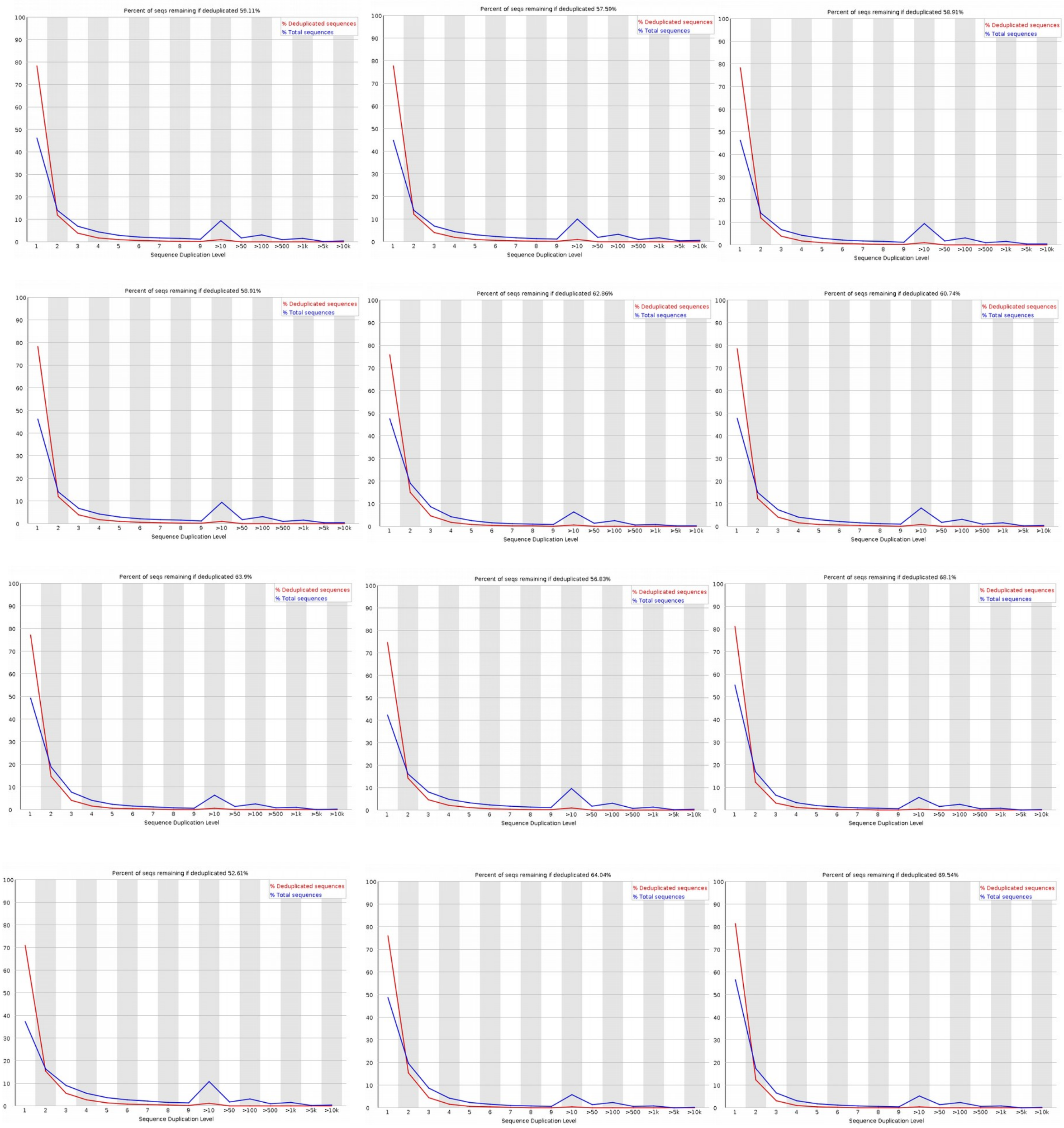

**Figure S2.** GC content of sample gDNA compared with theoretical normal distribution for each replicate performed using FastQC.

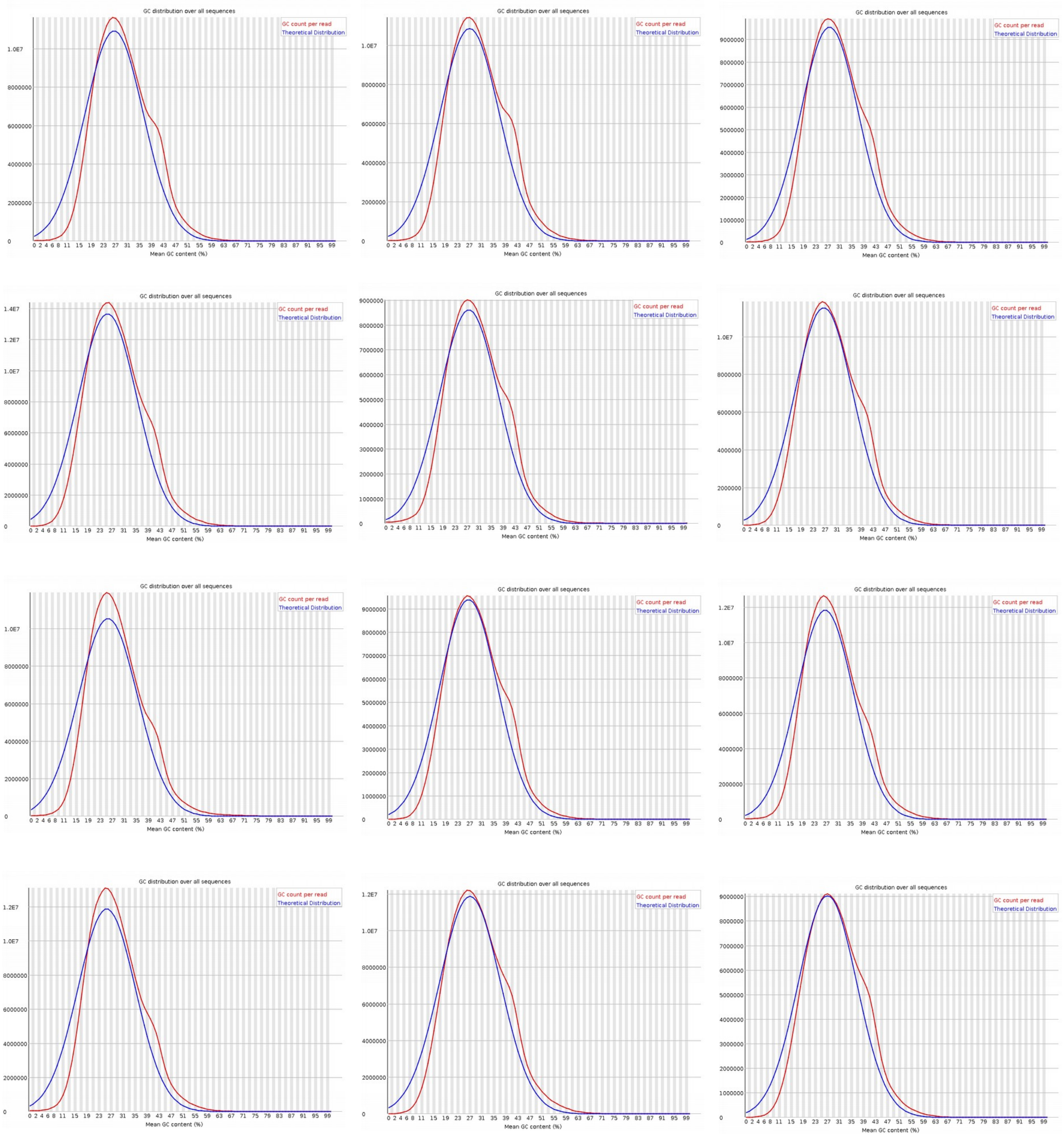

**Figure S3.** Histograms of sample coverage demonstrating the frequency of length of reads within each dose of ATZ in $log_{10}$ of read coverage per base.

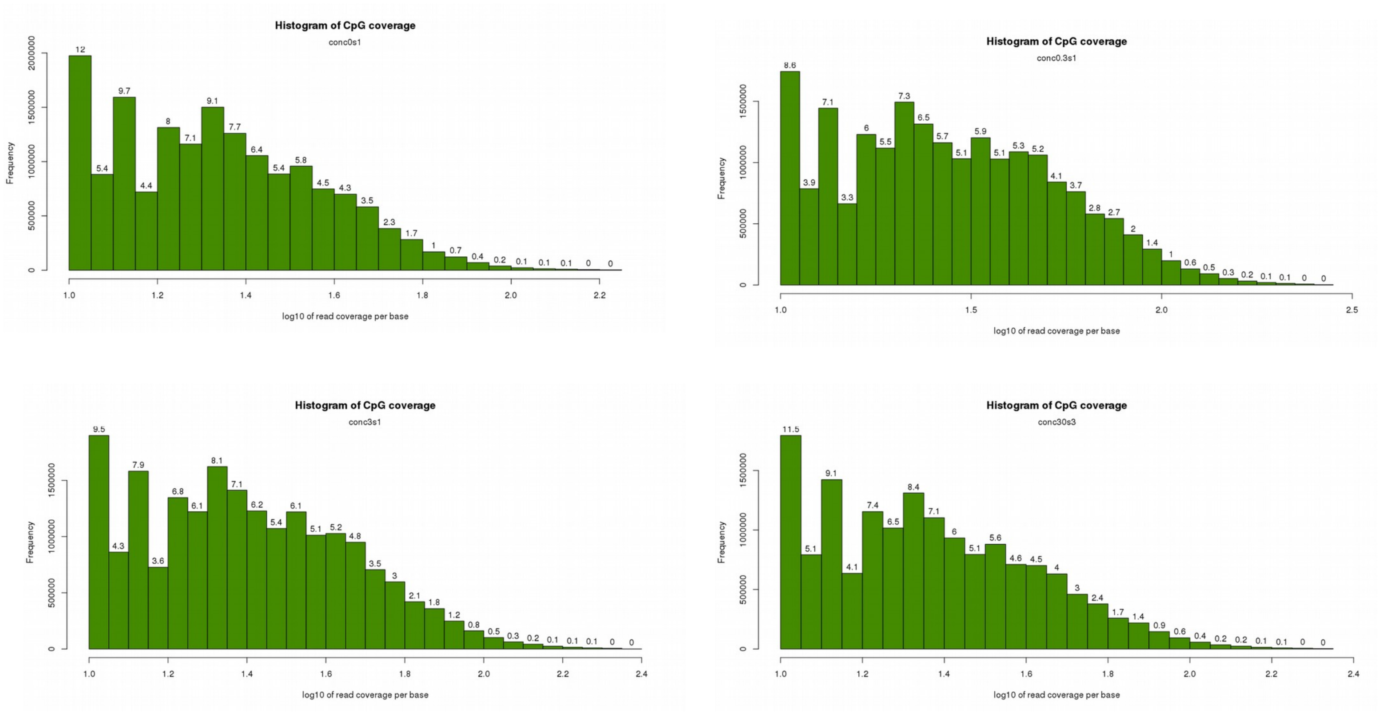

**Figure S4.** Heatmap comparing the average methylation percentage of all genes. Color scale represents the average proportion of methylated cytosines at CpG throughout the gene, with each bar representing a unique gene. From innermost to outermost ring: 0 ppb ATZ, 0.3 ppb ATZ, 3 ppb ATZ, 30 ppb ATZ.

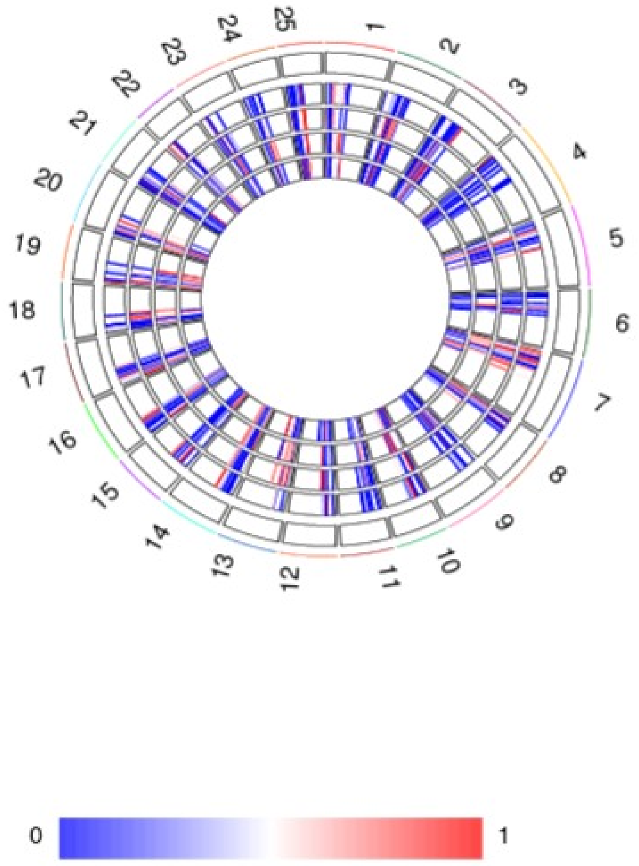

**Figure S5. A.** Number of DMR per chromosome in analysis comparing 0.3 ppb ATZ and 3 ppb ATZ. **B.** Number of DMR per chromosome in analysis comparing 0.3 ppb ATZ and 30 ppb ATZ **C.** Number of DMR per chromosome in analysis comparing 3 ppb ATZ and 30 ppb ATZ.

**A.** Comparison of samples treated with 0.3 ppb and 3 ppb ATZ

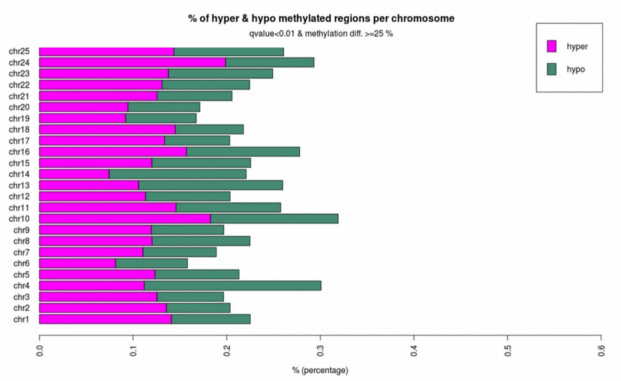


**B.** Comparison of samples treated with 0.3 ppb and 30 ppb ATZ

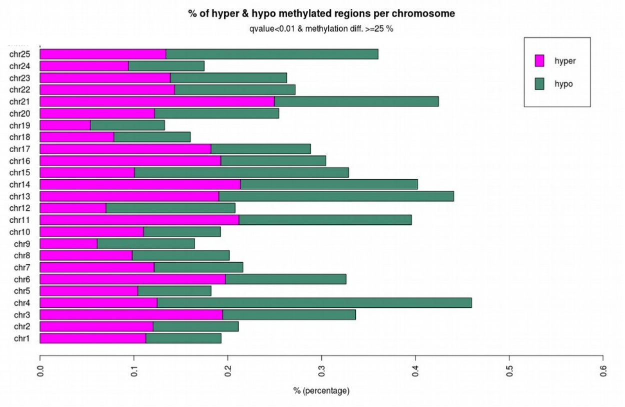


**C.** Comparison of samples treated with 3 ppb and 30 ppb ATZ

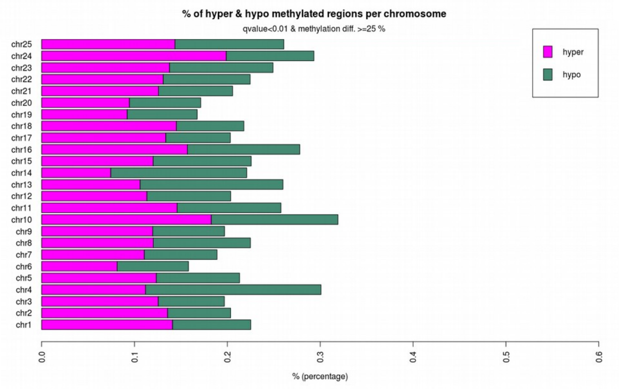

**Figure S6.** Proportion of DMR located in exons, introns, promoters (1000 kb upstream), and intergenic regions for 0 ATZ- 3 ppb ATZ comparison, 0 ATZ- 30 ppb ATZ comparison, 0.3 ATZ- 3 ppb ATZ comparison, 0.3ATZ- 30 ppb ATZ comparison, and 3 ATZ- 30 ppb ATZ comparison.

(0 ppb – 3 ppb) (0 ppb – 30 ppb)

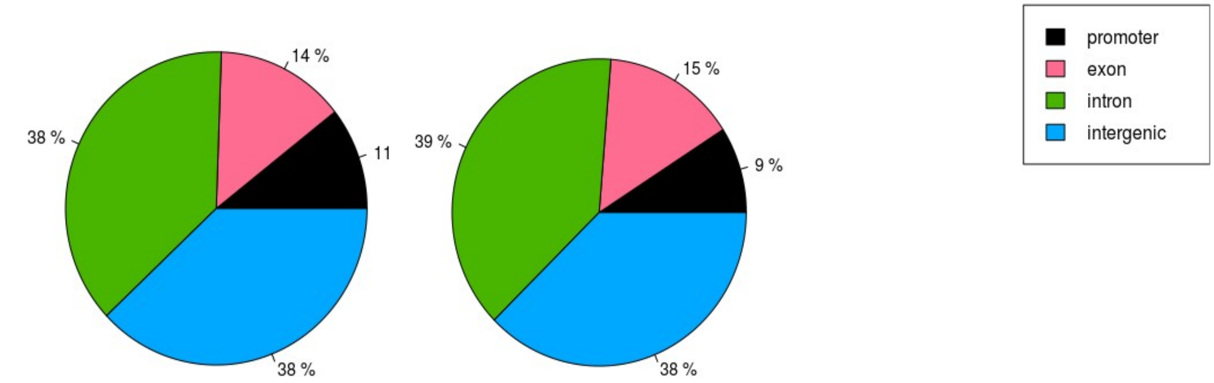


(0.3 ppb – 3 ppb) (0.3 ppb – 30 ppb)

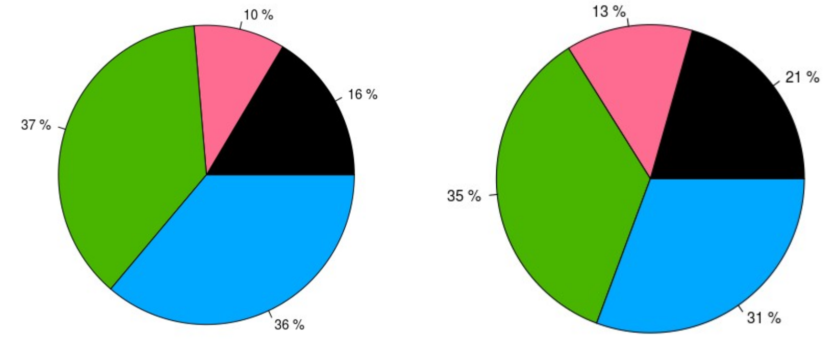


(3 ppb – 30 ppb)

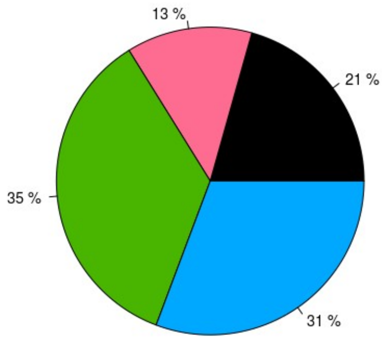

**Figure S7.** Proportion of DMR located in CpGi and CpG shores for 0 ATZ- 3 ppb ATZ comparison, 0 ATZ- 30 ppb ATZ comparison, 0.3 ATZ- 3 ppb ATZ comparison, 0.3 ATZ- 30 ppb ATZ comparison, and 3 ATZ- 30 ppb ATZ comparison.

(0 ppb – 0.3 ppb)

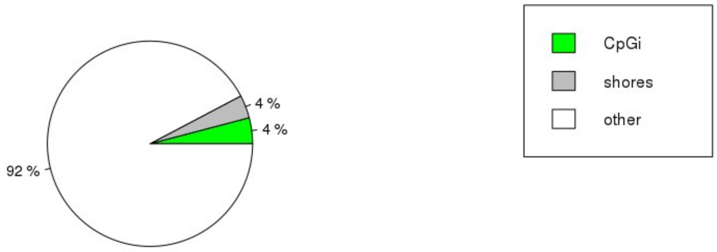


(0 ppb – 3 ppb)

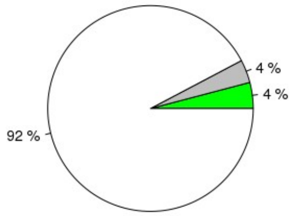


(0 ppb – 30 ppb)

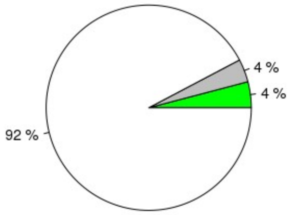


(3 ppb – 30 ppb)

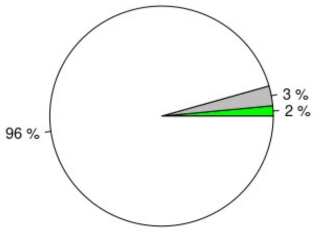

(0.3 ppb – 30 ppb)

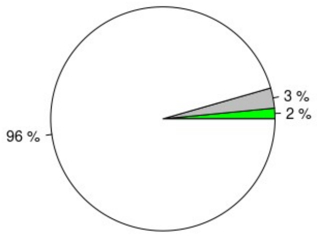


(0.3 ppb – 3 ppb)

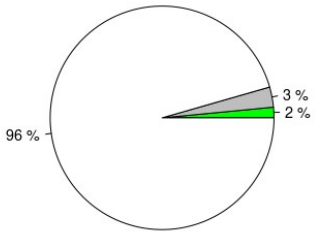

**Figure S8.** Average methylation percentage in DMGs containing CpG shores (-2000 to +2000 bp of TSS).

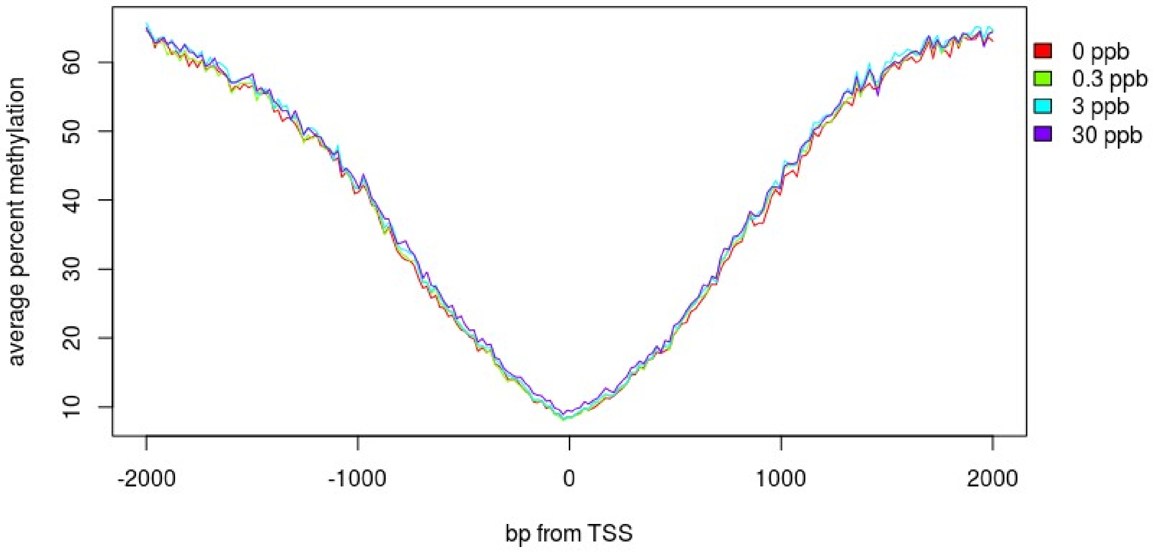

**Figure S9. A.** PCA of replicates using CpG methylation of promoter region, **B.** Hierarchal clustering of replicates using CpG methylation of promoter, **C.** PCA of replicates using CpG methylation of exon, **D**. Hierarchal clustering of replicates using CpG methylation of exon.

**A.**

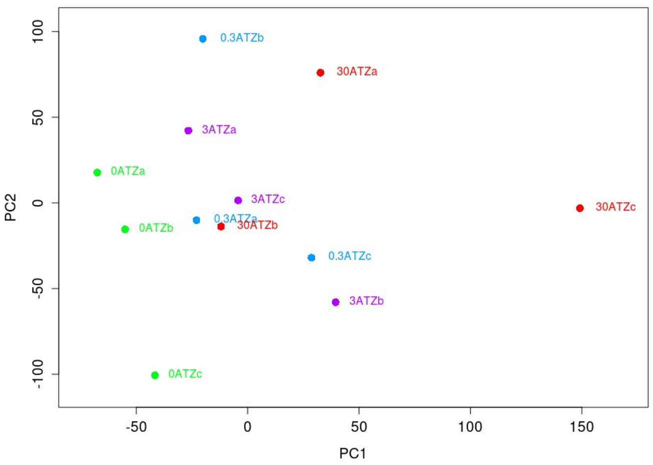


**B.**

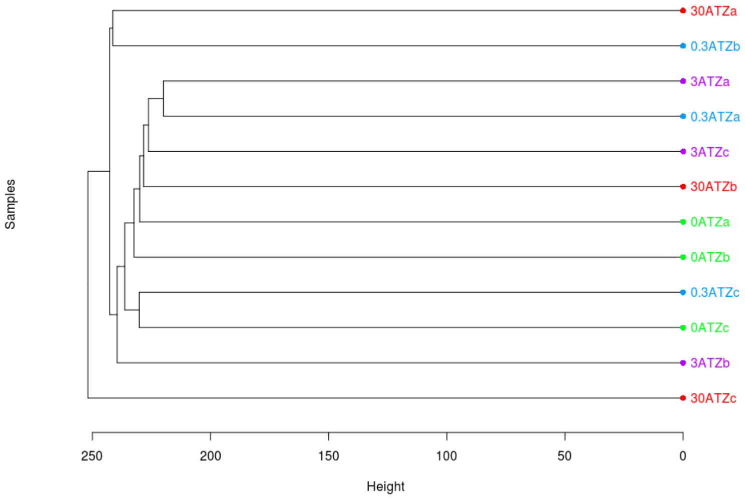

**C.**

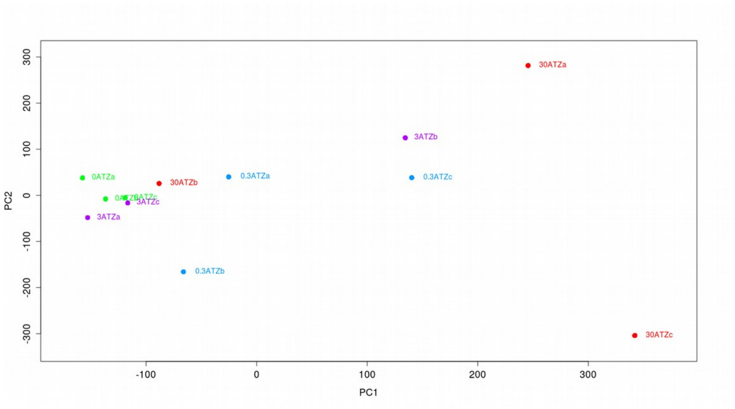
30ATZa
3ATZb
0ATZa
30ATZb
0.3ATZa
0.3ATZc
3ATZc
3ATZa
0.3ATZb
30ATZc
PC2
PC1
300
200
100
0
-100
-200
-300
-100
0
100
200
300

**D.**

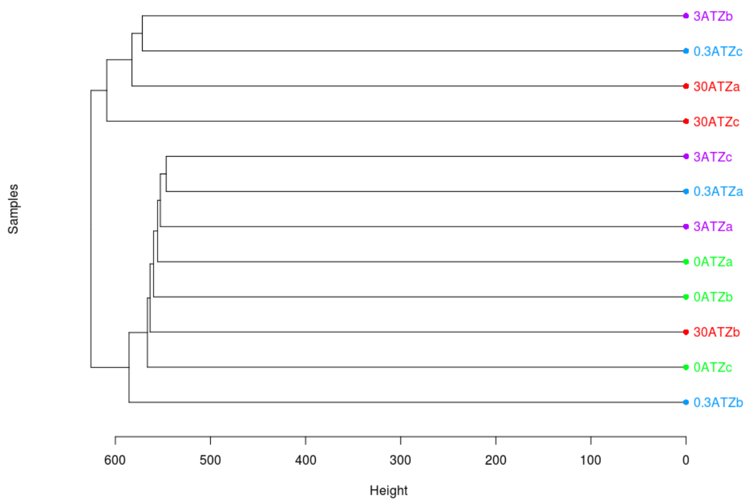
3ATZb
0.3ATZc
30ATZa
30ATZc
3ATZc
0.3ATZa
3ATZa
0ATZa
0ATZb
30ATZb
0ATZc
0.3ATZb
Samples
600
500
400
300
200
100
0
Height

**Figure S10. A.** Venn diagram demonstrating the number of genes with differentially methylated promoter in each comparison group. **B**. Venn diagram demonstrating the number of genes with differentially methylated gene body in each comparison group.

**A.**

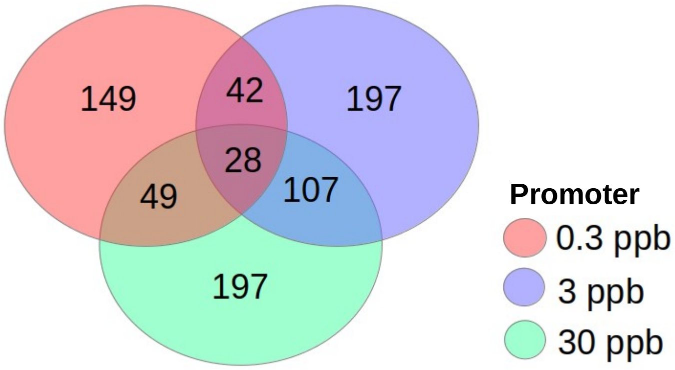


**B.**

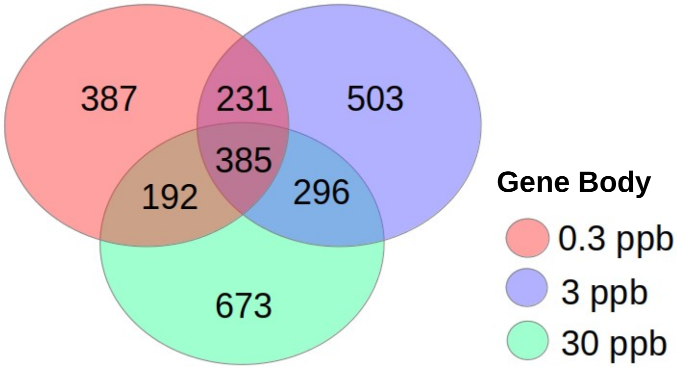

**Figure S11. A - C**. DMG from 0 ppb – 3 ppb comparison (marked in purple) within the GABA Receptor signaling pathway (**A**), Long-Term Synaptic Depression pathway (**B**), and Gonadotropin-Releasing Hormone Signaling pathway (**C**).

**A.** GABA Receptor Signaling

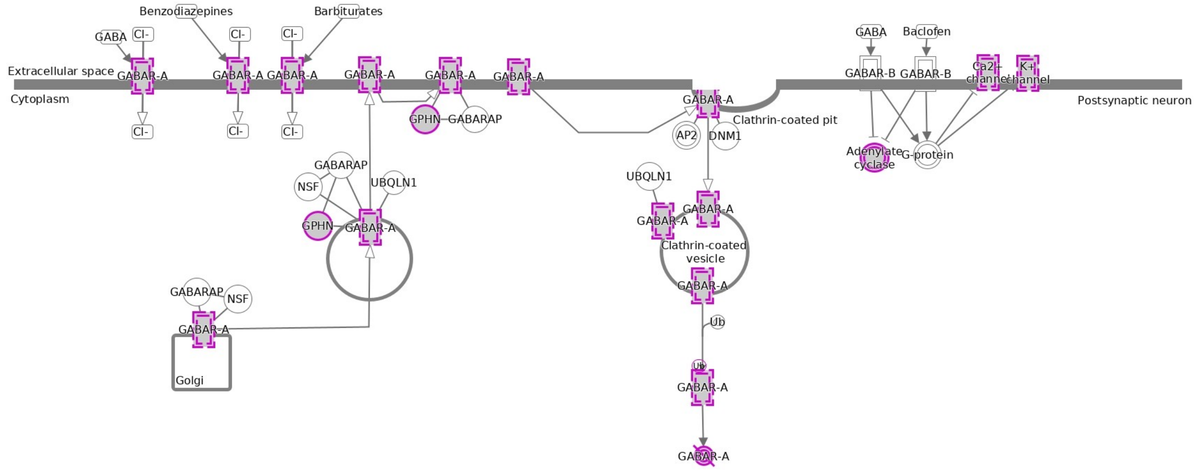


**B.** Long-Term Synaptic Depression

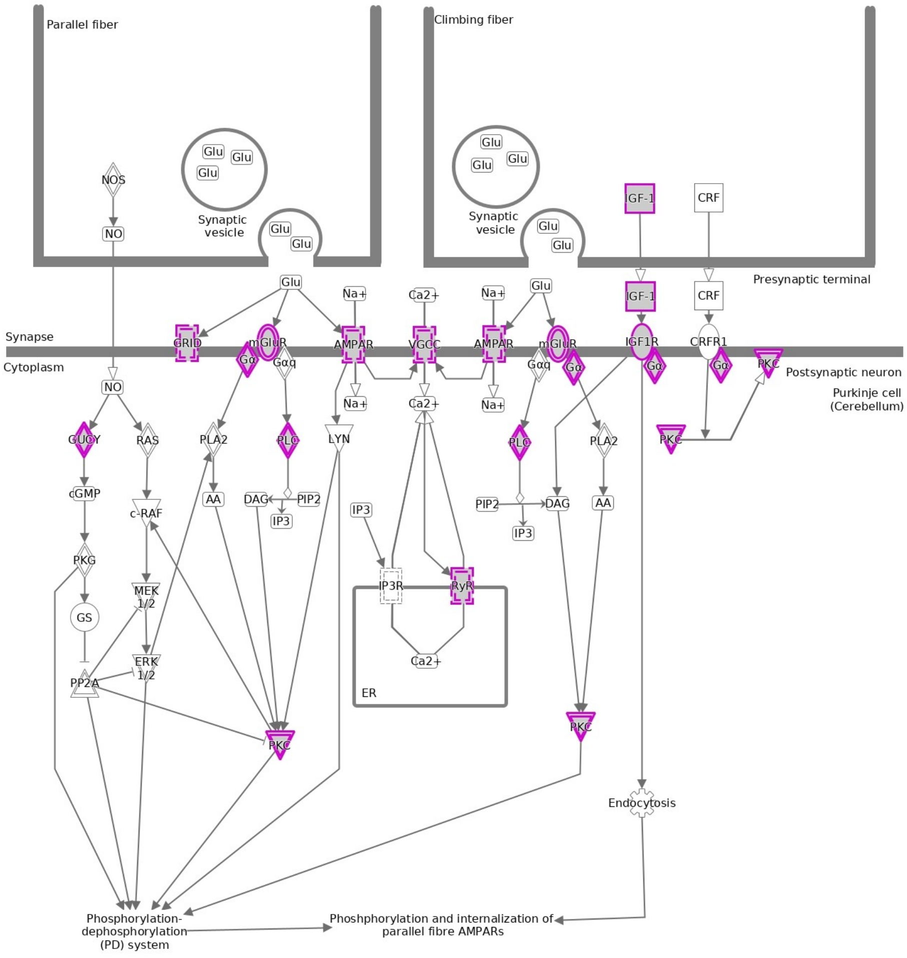

**C.** Gonadotropin-Releasing Hormone Signaling

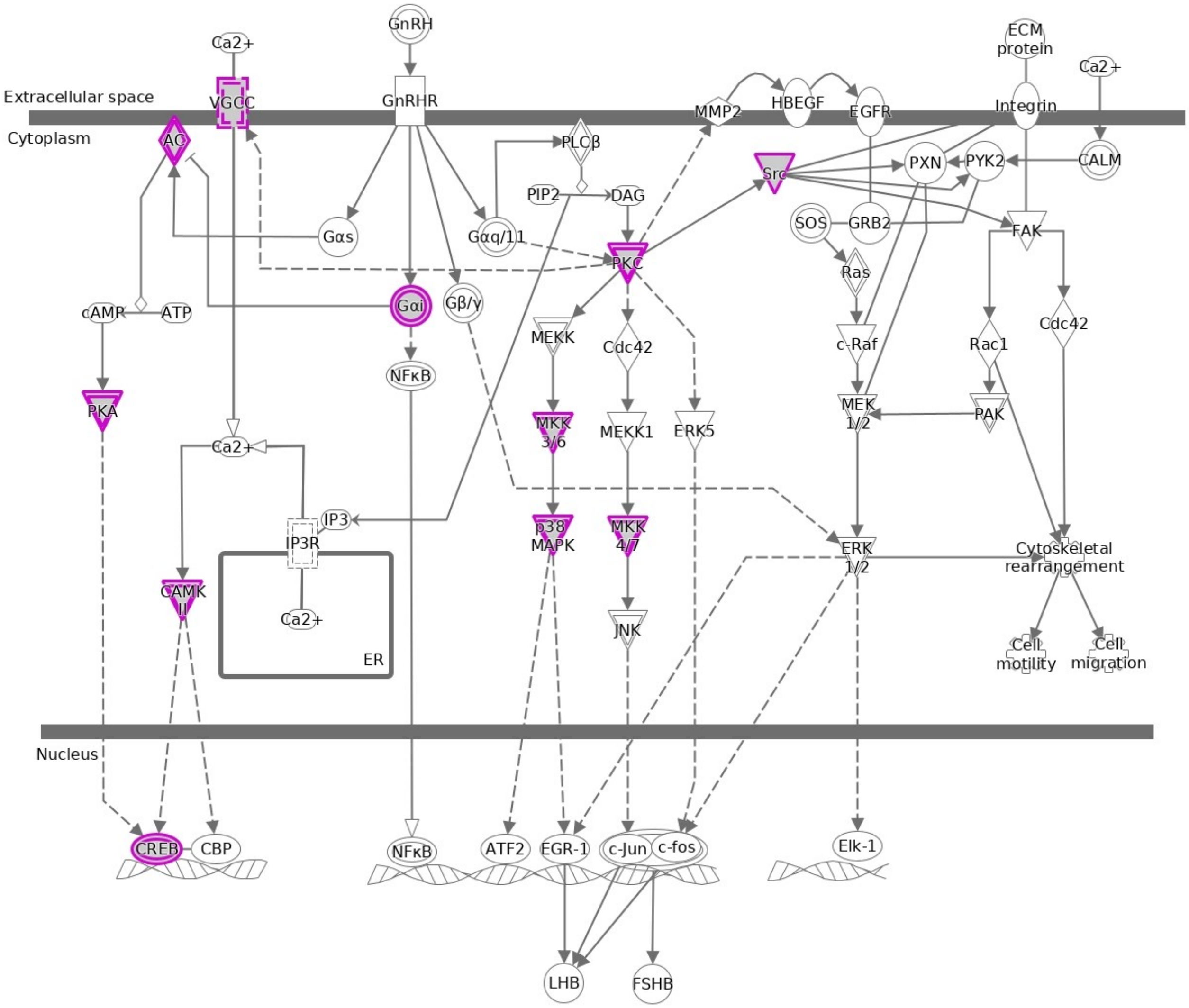

**Figure S12. A – C.** Scatterplots of differentially methylated and expressed genes with methylation difference on x-axis and expression level fold change on y-axis for genes containing differentially methylated exons identified in (**A**) 0 - 0.3 ppb, (**B**) 0 - 3 ppb and (**C**) and 0 - 30 ppb comparisons. Similar plots were made for common genes with differentially methylated intron sequences as shown for genes identified in (**D**) 0 - 0.3 ppb, (**E**), 0 - 3 ppb and (**F**) and 0 - 30 ppb comparisons.

**A.**
0.3 ppb exons

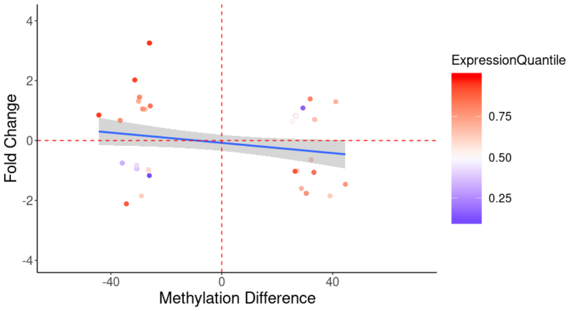


**B.**
3 ppb exons

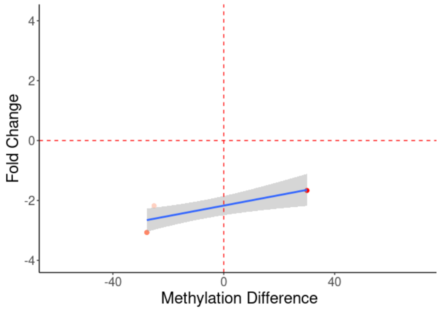


**C.**
30 ppb exon

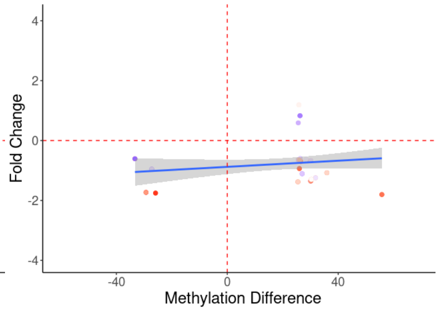


**D.**
0.3 ppb introns

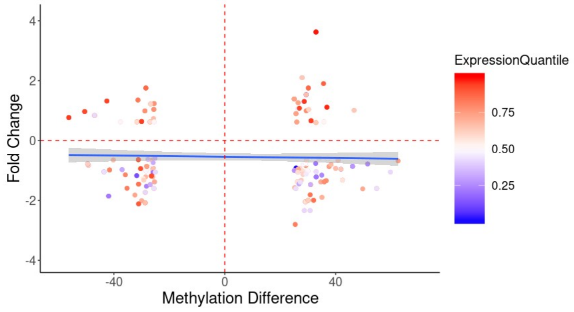


**E.**
3 ppb introns

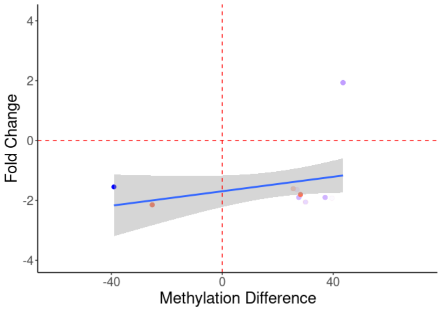


**F.**
30 ppb introns

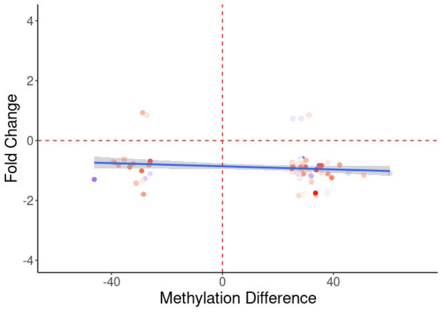

**Figure S13.** Illustration of procedure for husbandry and treatment of zebrafish embryos.

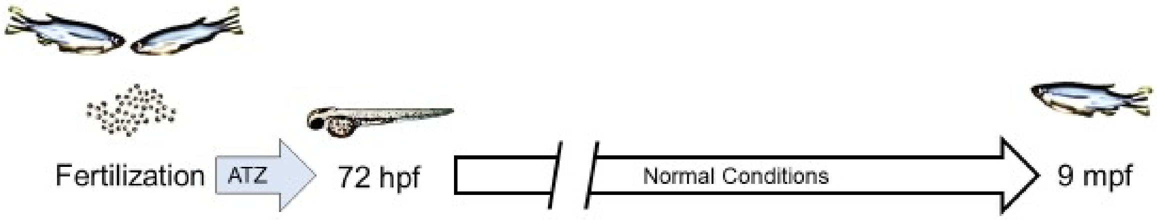
